# 3D Mapping of Neurofibrillary Tangle Burden in the Human Medial Temporal Lobe

**DOI:** 10.1101/2021.01.15.421909

**Authors:** Paul A. Yushkevich, Mónica Muñoz López, Maria Mercedes Iñiguez de Onzoño Martin, Ranjit Ittyerah, Sydney Lim, Sadhana Ravikumar, Madigan L. Bedard, Stephen Pickup, Weixia Liu, Jiancong Wang, Ling Yu Hung, Jade Lasserve, Nicolas Vergnet, Long Xie, Mengjin Dong, Salena Cui, Lauren McCollum, John L. Robinson, Theresa Schuck, Robin de Flores, Murray Grossman, M. Dylan Tisdall, Karthik Prabhakaran, Gabor Mizsei, Sandhitsu R. Das, Emilio Artacho-Pérula, María del Mar Arroyo Jiménez, María Pilar Marcos Rabal, Francisco Javier Molina Romero, Sandra Cebada Sánchez, José Carlos Delgado González, Carlos de la Rosa-Prieto, Marta Córcoles Parada, Edward B. Lee, John Q. Trojanowski, Daniel T. Ohm, Laura E.M. Wisse, David A. Wolk, David J. Irwin, Ricardo Insausti

**Affiliations:** Department of Radiology, University of Pennsylvania, Philadelphia, USA; Human Neuroanatomy Laboratory, Neuromax CSIC Associated Unit, University of Castilla-La Mancha, Albacete, Spain; Department of Neurology, University of Pennsylvania, Philadelphia, USA; Department of Pathology, University of Pennsylvania, Philadelphia, USA; Institut National de la Santé et de la Recherche Médicale (INSERM), Caen, France; Department of Diagnostic Radiology, University of Lund, Lund, Sweden

## Abstract

Tau protein neurofibrillary tangles (NFT) are closely linked to neuronal/synaptic loss and cognitive decline in Alzheimer’s disease (AD) and related dementias. Our knowledge of the pattern of NFT progression in the human brain, critical to the development of imaging biomarkers and interpretation of in vivo imaging studies in AD, is based on conventional 2D histology studies that only sample the brain sparsely. To address this limitation, ex vivo MRI and dense serial histological imaging in 18 human medial temporal lobe (MTL) specimens were used to construct 3D quantitative maps of NFT burden in the MTL at individual and group levels. These maps reveal significant variation in NFT burden along the anterior-posterior axis. While early NFT pathology is thought to be confined to the transentorhinal region, we find similar levels of NFT burden in this region and other MTL subregions, including amygdala, temporopolar cortex, and subiculum/CA1.

## 1. Introduction

Tau neurofibrillary tangle (NFT) pathology is closely linked to neurodegeneration and cognitive decline in Alzheimer’s disease (AD) [13]. The topographic characterization of the spread of NFT pathology through the human brain by Braak and others [15, 47, 5, 16, 6, 17, 18, 19, 36] has had tremendous impact in many areas of AD research, including the diagnosis of AD [36, 38]. Yet this characterization is based on histological examination of postmortem tissue that is inherently two-dimensional (2D) and samples the brain at a sparse set of locations, limiting our knowledge of the spread of NFT pathology. In particular, since histological sectioning is almost always done in the coronal plane, little is known about the distribution of NFT pathology along the anterior-posterior axis.

Recent advances in positron emission tomography (PET) imaging enabled in vivo detection and mapping of tau pathology in three dimensions (3D). Tau PET imaging suggests a more diffuse pattern of early spread of tau pathology [39] than what is suggested by the Braak and Braak staging system [16, 17, 36], according to which early NFT pathology is largely confined to the transentorhinal cortex, a small region located on the medial portion of Brodmann Area 35 (BA35) in the anterior medial temporal lobe (MTL). However, tau PET is not a direct measure of NFT burden in the brain, since it has limited spatial resolution and has variable binding to multiple types of tau pathology. Structural magnetic resonance imaging (MRI) studies examining patterns of neurodegeneration in the MTL also suggest that, while BA35 is clearly impacted early in the disease, other MTL structures undergo similar rates of atrophy [77, 79]. The analysis of in vivo PET and MRI is significantly hampered by the lack of a comprehensive “gold standard” postmortem reference that would characterize the distribution and spread of NFT pathology in 3D and would be compatible with tools used analyze in vivo PET and MRI data.

Quantitative 3D mapping of NFT and other neurodegenerative proteinopathies may also hold the key to solving the problem of mixed pathology in Alzheimer’s disease (AD). Most patients diagnosed with AD at autopsy also harbor one or more concomitant neurodegenerative pathologies, such as TDP-43 and α-synuclein proteinopathies, non-AD tauopathies, as well as vascular disease [63, 36, 70, 7, 60, 75, 40, 55]. Unlike β-amyloid and tau, these pathologies cannot be detected reliably in vivo using current technology, although efforts are underway to develop PET tracers that would allow their detection. Histological staging studies and antemortem-pathology correlation studies suggest that these concomitant pathologies follow a pattern of spread that is different from NFT pathology, leading to distinct patterns of brain tissue loss and atrophy [29, 24, 21, 41, 10, 55, 28]. Precise 3D characterization of the spread of NFT pathology, contrasted to 3D characterization of the spread of concomitant pathologies may help define “hot spots” in the brain where atrophy is more likely to be linked to one pathology than another, potentially allowing in vivo detection. Furthermore, such hot spots could help clinical trials measure treatment efficacy more reliably by quantifying the brain’s response to treatment in relevant hot spots, rather than less specific regions such as the hippocampus. Additionally, tools that allow 3D mapping of TDP-43 and α-synuclein proteinopathies could prove useful for validation of novel PET tracers.

In this paper, we develop a framework that can generate quantitative 3D maps of neurodegenerative proteinopathies in the human brain, which can then serve as a 3D reference for in vivo MRI analysis. We restrict our attention to the medial temporal lobe (MTL), an essential component of the human memory system and the site of early neurodegeneration in AD. Tau, TDP-43, α-synuclein and vascular pathologies all affect the MTL in their early stages, making the MTL a hotbed of early neurodegenerative activity [66, 36, 55]. We also restrict our attention to generating 3D maps of tau NFT burden, although the underlying approach is amendable to similar mapping of other pathologies. Our framework combines ex vivo MRI, image-guided tissue processing, serial histology imaging, and advanced machine learning algorithms. We leverage it to generate 3D maps of NFT burden for 18 brain donors, culminating in group-level maps that reveal significant early involvement of MTL structures beyond the transentorhinal cortex, as well as a marked anterior to posterior gradient of NFT deposition. These 3D maps of NFT deposition, defined in the space of an *in vivo* brain MRI template, are provided in digital form to facilitate their use in in vivo MRI and PET analysis.

## 2. Results

Brain hemisphere specimens from 18 donors 45-93 years of age were obtained from the archive cases from the Human Neuroanatomy Laboratory at the University of Castilla La Mancha (UCLM, n=12) and the Center for Neurodegenerative Disease Research at the University of Pennsylvania (UPenn, n=6). In each donation, the donor’s next of kin provided consent to autopsy. Donors from UCLM were from the general population served by the brain bank, and included mostly older adults with no known neurological disease. Donors from UPenn were participants in *in vivo* aging and dementia research, and included patients from the Penn Frontotemporal Degeneration Center and the Penn Alzheimer’s Disease Core Center. Table 1 provides summary demographic and diagnostic data for the brain donor cohort, with additional details in Supplemental Table S.1.

**Table 1:** Demographic composition of the brain donor cohort, primary and secondary postmortem diagnoses, and global neuropathological staging using the Hyman et al. [36] protocol. Staging and diagnoses were derived from the opposite hemisphere from the one scanned with MRI. (*): For the brain donor with the diagnosis of argyrophilic grain disease, the Braak (B) score was undeterminable.

| Brain Donor Cohort |  |  |  |  |  |
| --- | --- | --- | --- | --- | --- |
| N |  | 18 |  |  |  |
| Age |  | 75.2±11.4 | 45 - 93 |  |  |
| Sex |  | 7F, 11M |  |  |  |
| Diagnosis (from contralateral sampling) |  | Primary | Secondary |  |  |
|  | Unremarkable brain | 1 |  |  |  |
|  | Primary age-related pathology (PART) | 4 |  |  |  |
|  | Low AD neuropathological change (ADNC) | 6 | 1 |  |  |
|  | Intermediate ADNC | 1 |  |  |  |
|  | Corticobasal Degeneration (CBD) | 2 | 1 |  |  |
|  | Lewy Body Disease (LBD) | 1 | 2 |  |  |
|  | Argyrophilic Grain Disease (AGD) | 1 |  |  |  |
|  | Frontotemporal Lobar Degeneration (TDP-43 type) | 1 | 1 |  |  |
|  | Cerebrovascular Disease (CVD) | 1 | 1 |  |  |
|  | Progressive Supranuclear Palsy (PSP) |  | 1 |  |  |
| Neuropathological Staging<br>(from contralateral sampling) |  | Stage |  |  |  |
|  |  | 0 | 1 | 2 | 3 |
|  | Amyloid (A) | 7 | 10 | 0 | 1 |
|  | Braak (B)* | 3 | 10 | 4 | 0 |
|  | CERAD (C) | 13 | 4 | 0 | 1 |
| | α-synuclein | 15 | 2 | 1 | 0 |
|  | TDP-43 | 16 | 0 | 0 | 2 |

For each brain donation, a series of imaging procedures summarized in Figure 1 were performed. Tissue from one brain hemisphere was used for imaging (MRI and histology) and the opposite hemisphere was sampled for diagnostic pathology following the NIA-AA protocol [36]. After fixation, a specimen containing the intact MTL was dissected from the hemisphere. Each intact MTL specimen was scanned overnight on a 9.4T animal MRI scanner with 0.2 x 0.2 x 0.2 mm^3^ native resolution using a standard T2-weighted sequence. A separate T2-weighted MRI scan was obtained on a 7 Tesla human MRI scanner with 0.4 x 0.4 x 0.4 mm^3^ resolution. Registration between the 9.4T and 7T scans was performed to correct the 9.4T scan for geometric distortions due to the non-linearity of the magnetic gradient field that increases towards the ends of the sample. A custom mold that tightly fits the MTL specimen was 3D printed using the 7T scan and used to guide tissue sectioning. The mold (Figure 1) orients sectioning orthogonal to the main axis of the hippocampus and reduces the complexity of subsequent registration between MRI and histology, as the plane of histological sectioning relative to the 7T MRI scan is known. Using the custom mold, specimens were cut into 20 mm thick blocks, with most specimens yielding 4 blocks. Cryoprotected blocks were frozen using dry ice and sectioned using a sliding microtome coupled to a freezing unit into 50 *μ*m sections, with no gaps between sections. Before cutting each section, a digital photograph of the block were taken using a mounted overhead camera (called *blockface images*). Every tenth section (sections 10, 20, 30,…) was stained for the Nissl series using the thionin stain. Every twentieth section (19, 39, 59,…) was stained using AT8, a human phosphorylated tau antibody IHC stain, and were counterstained for Nissl. Thus, Nissl stained sections were at 0.5mm intervals (^~^40 per block) and anti-tau sections were adjacent to the Nissl sections and at 1mm intervals (^~^20 per block). Sections were mounted on 75mm x 50mm glass slides, digitally scanned at 20X resolution, and uploaded to an in-house created cloud-based digital histology archive that supports web-based visualization, anatomical labeling, and machine learning classifier training.

**Figure 1:**
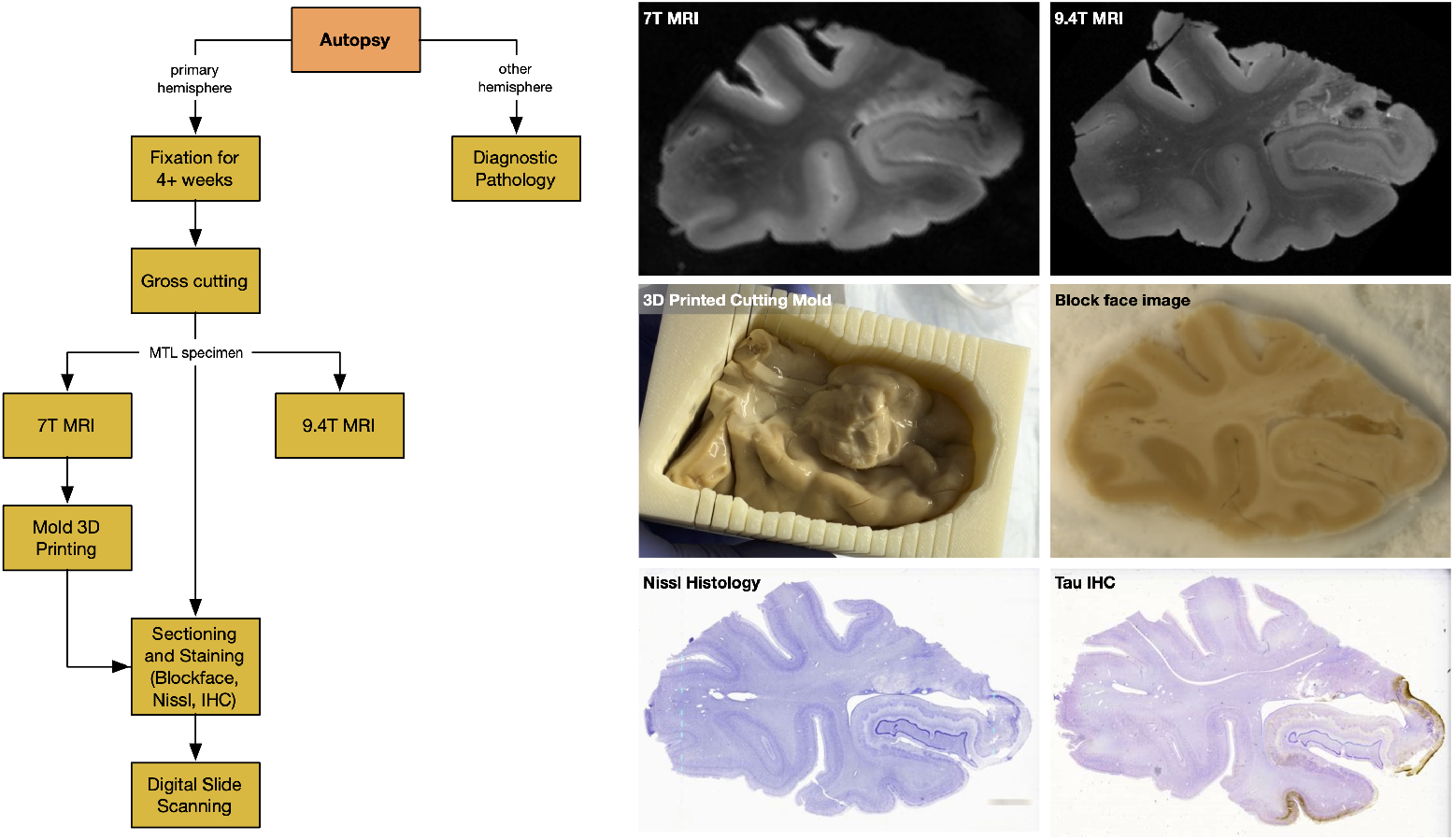
The workflow of specimen preparation and imaging, along with examples of different imaging modalities and a sample 3D printed mold. The six images show a coronal section from the 0.4 × 0.4 × 0.4 mm^3^ 7T MRI, which is used to generate 3D printed molds and to correct 9.4T scans for distortion; a coronal section from the 0.2 × 0.2 × 0.2 mm^3^ 9.4T MRI; intact MTL specimen placed in the cutting mold 3D printed from the 7T MRI; a blockface photograph taken during cryosectioning; a Nissl-stained histology slide; and an anti-tau IHC slide counterstained for Nissl. Abbreviations: MRI: magnetic resonance imaging; IHC: immunohistochemistry; T: Tesla.

### 2.1. Automated Measures of Tau NFT Burden Agree with Conventional Histopathology Measures

#### 2.1.1. Maps of NFT Burden Generated from Whole-Slide Images using Weakly Supervised Learning

The first stage of our framework was to generate “heat maps” that quantify the burden of NFT pathology on individual anti-tau IHC sections using deep learning. We adapted the *weakly supervised learning (WSL) approach* from computer vision, which performs segmentation at the level of pixels (e.g., outlines animals in photos) using training data that contains only image-level labels (e.g., “image *I* contains a cat”). WSL occupies the ideal middle ground between simply classifying image regions into tangle/non-tangle classes (which fails to discriminate between regions with dense tangles and sparse tangles) and full-blown segmentation of individual tangles (which requires costly pixel-level training data, and is unnecessary when the objective is to derive NFT burden maps at the resolution of the ex vivo MRI).

Training data for WSL were generated by 12 raters using a custom web-based slide annotation system over the course of two day-long “Tanglethon” events. Over 11,000 512×512 pixel patches (examples in Supplemental Figure S.2) were extracted from 176 slides in six MTL specimens and assigned into tangle-like (NFTs and pre-tangles) and non-tangle (tau neuropil threads, astroglial tau, tau coils in the white matter, normal tissue, slide background, artifacts, tissue folds) classes. The *WildCat* WSL algorithm [31] was trained and evaluated in a leave-one-out cross-validation setting, using slides from five specimens for training/validation and using the remaining specimen for testing. WildCat assigned testing patches to the correct class (tangle or non-tangle) with the accuracy if 95.9 ± 2.0% (range 93.1% to 98.4%) across the six cross-validation experiments. For each input patch, WildCat yielded a heat map indicating the location and intensity of the tangles (examples in Supplemental Figure S.2).

#### 2.1.2. Automated NFT Burden Measure is Concordant with Manual Counting

A single WildCat model was trained on all six Tanglethon specimens and applied to all anti-tau wholeslide IHC images in the study, yielding whole-slide NFT burden maps, illustrated in Supplemental Figure S.3. To validate these burden maps, 48 boxes of 2048 × 2048 pixels were sampled from three specimens not used for WildCat training, and tangle-like inclusions were counted in each box. The correlation between WildCat-derived NFT burden measure integrated over each box and the manual tangle count, plotted in Fig. 2a, was high (Kendall’s *τ* = 0.81, Spearman’s *ρ* = 0.94). Examining the boxes corresponding to points with high residual values in this plot suggests that WildCat-reported burden is greater than expected for a given number of tangles per box when the tangles are large and prominent (point II in 2), and less than expected when pre-tangles are present (point III in 2). This suggests that the WildCat NFT burden measure captures both the number and prominence of tangle-like pathological inclusions.

**Figure 2:**
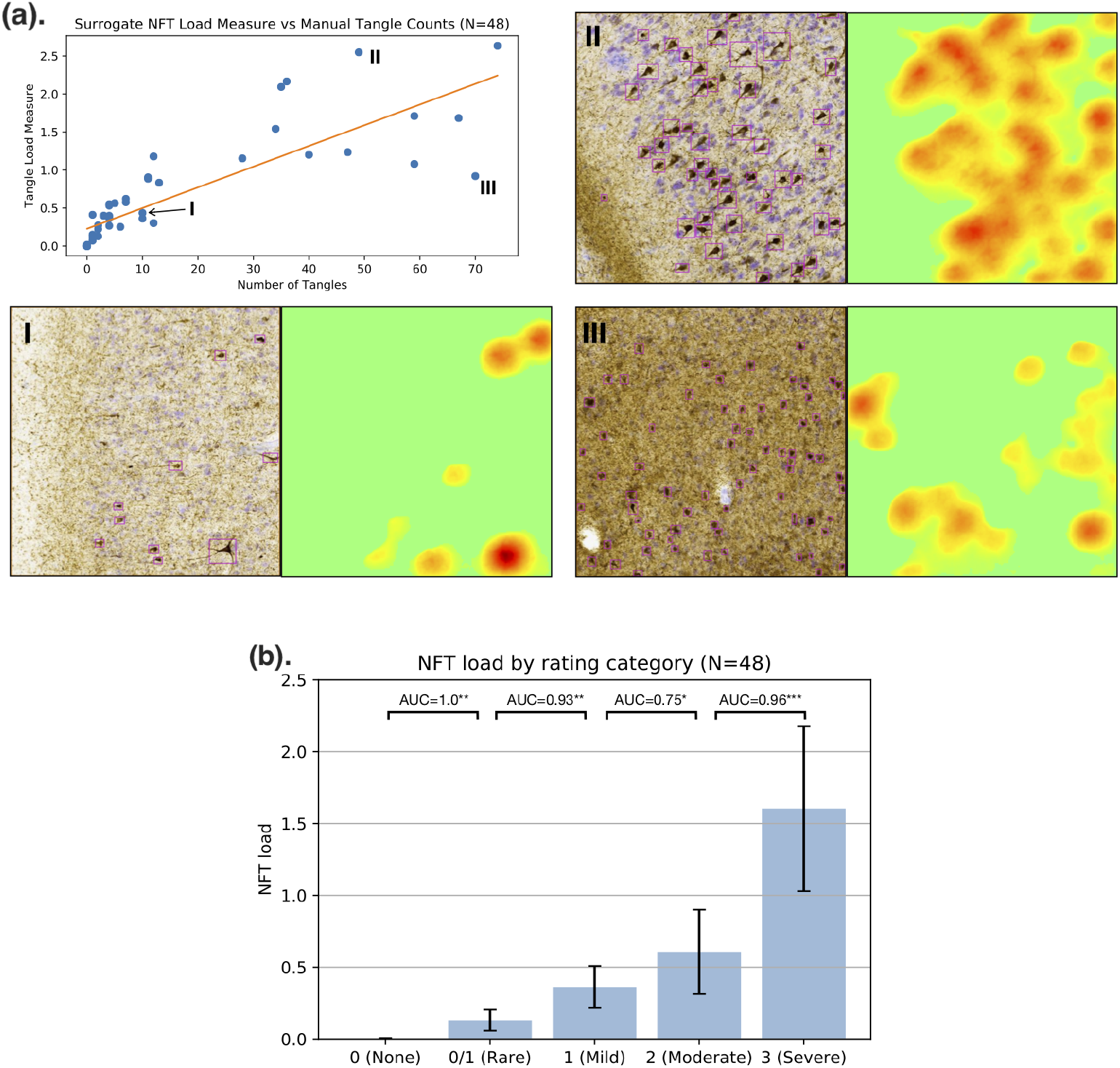
NFT burden measure derived using the weakly supervised learning algorithm WildCat compared to manual counting of NFTs and semi-quantitative ordinal ratings. (a). Regression between the WildCat burden measure and manual NFT count in 48 boxes from specimens left out from WildCat training is plotted in the top left panel. For three boxes, including two outliers, histology (with counted NFTs outlined in purple) and WildCat-derived burden maps are shown. (b). Plot of the average WildCat burden measure across five NFT severity categories into which the 48 boxes were assigned by an expert (DJI) based on visual rating. Mann-Whitney AUC (*U*/(*n*_1_*n*_2_)) is shown between adjacent categories (*: *p* < 0.05; **: *p* < 0.01; ***:p < 0.001, uncorrected)

#### 2.1.3. Automated NFT Burden Measure is Concordant with Semi-Quantitative Scores

Each of the 48 boxes was also assigned an ordinal semi-quantitative rating of NFT pathology by an expert (DJI), with categories ranging from 0 (None) to 3 (Severe). The distribution of WildCat-derived NFT burden was compared between rating categories (Fig. 2b) and found to be significantly different across all adjacent rating categories, with most Mann-Whitney AUCs between adjacent categories exceeding 0.9 (except AUC=0.75 between “mild” and “moderate” categories). The discrimination between categories “None” and “Rare” is particularly strong, indicating that this burden measure is sensitive to early NFT pathology. Overall, the strong discrimination between clinical categories and the strong concordance of NFT burden with manual counts suggest that our automated surrogate measure of NFT load may be used as a stand-in for these manual measures, especially for the purpose of differentiating between regions of low and high NFT density.

### 2.2. 3D Reconstruction of Serial Histology Guided by 3D Printed Molds Matches Anatomical Features between MRI and Histology

For each specimen, Nissl and anti-tau IHC histology images were reconstructed in the space of the 9.4T MRI scan using a multi-stage pipeline that incorporates multiple image registration steps and takes advantage of the known orientation of the histology slicing plane with respect to the MRI. The pipeline, detailed in Section 5.4, is largely automated, with a small number of manual initialization steps that together require under two hours per specimen. An example reconstruction is illustrated in Figure 3.

**Figure 3:**
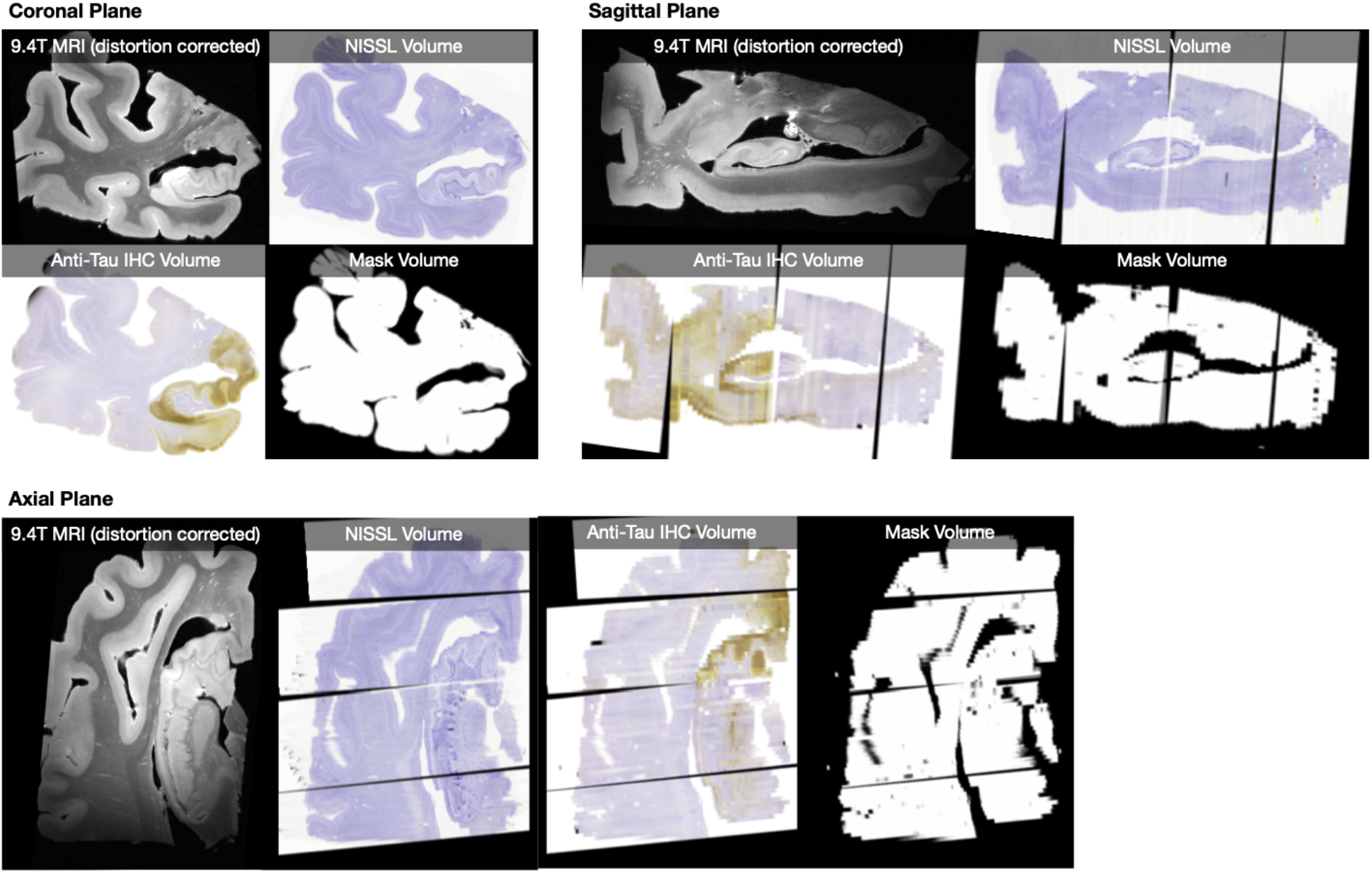
Example reconstruction, for a single specimen, of serial Nissl histology (one slide every 1mm) and serial anti-tau IHC histology (one slide every 2mm) in the space of the distortion-corrected 9.4T MRI. Reconstructions for all 18 specimens are included in the digital archive. The mask volume specifies where in the MRI image space histology measures are available.

To evaluate reconstruction accuracy, four anatomical curves were traced by the same rater (SAL) independently in Nissl and MRI space (Figure 4). Symmetric root mean squared distance (RMSD) between the curves was measured at different stages of 3D reconstruction, with larger values of distance indicating greater mismatch. At the final reconstruction stage, the mean RMSD between Nissl and MRI curves was 0.37±0.14mm for curve C1, 0.29±0.24mm for curve C2, 0.21 ±0.15mm for curve C3, and 0.37±0.29mm for curve C4. Examples of median registration performance for each curve are shown in Figure 4. Overall, the registration accuracy is high, with average RMSD below two MRI voxel widths for all four curves, and close to one voxel width for curves C2 and C3. Additional results describing curve mismatch at different stages of the registration pipeline are reported in Supplemental Section Appendix B.1.

**Figure 4:**
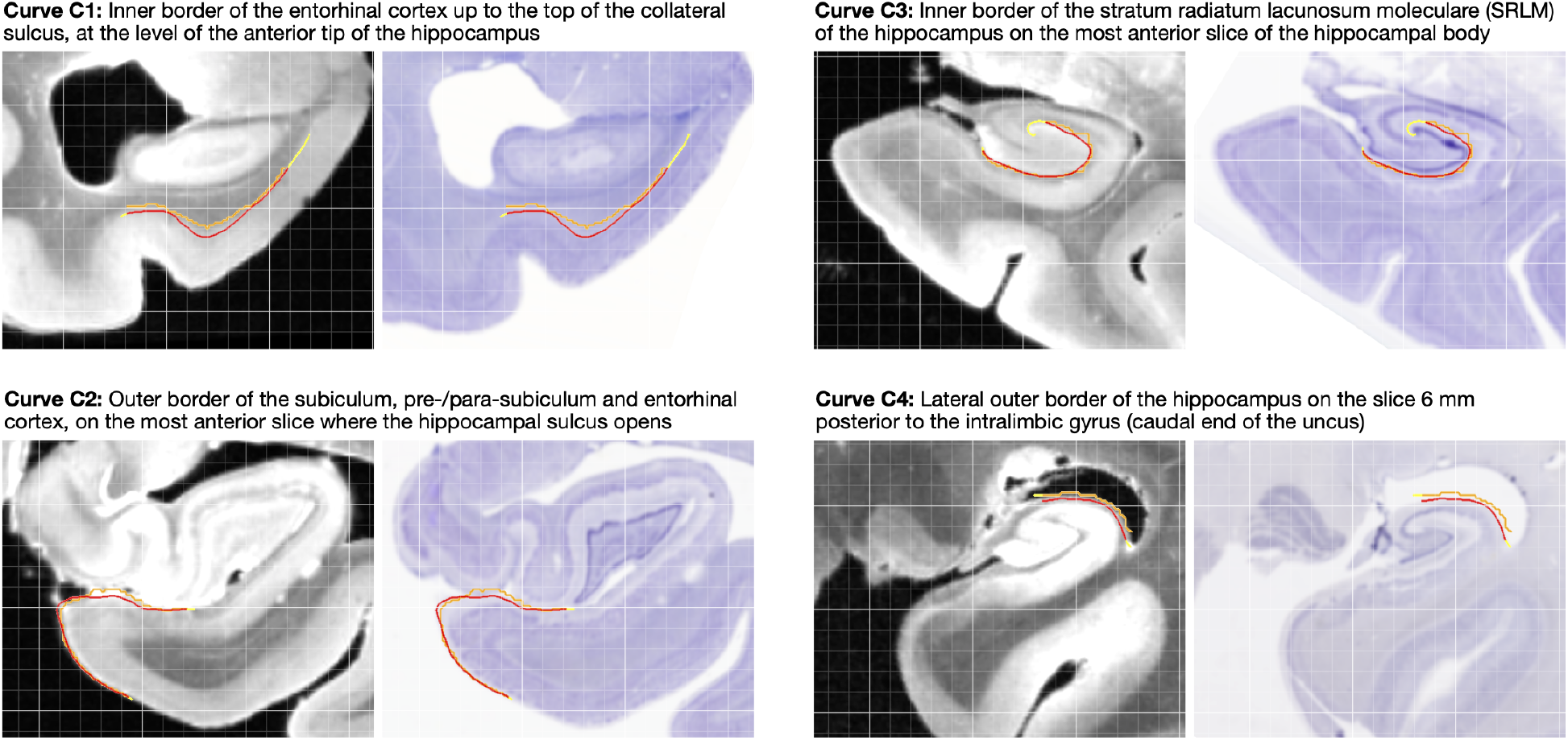
Examples of curve-based evaluation of registration between 9.4 Tesla MRI and reconstructed Nissl slides. Four anatomical curves (C1-C4) were traced independently in MRI (red) and Nissl space (orange). After registration, the symmetric root mean squared distance between the registered curves was measured, with yellow portions of the curves excluded. The four examples shown here correspond to median registration performance for each curve.

### 2.3. 3D Maps of NFT Density Generated in 18 MTL Specimens

Following 3D reconstruction, whole-slide maps of NFT burden generated by WildCat were transformed into the space of the distortion-corrected 9.4T MRI for all 18 specimens. Coronal and sagittal sections through these reconstructed 3D NFT burden maps are shown in Figure 5, and the full 3D maps are distributed in the digital archive. While some small errors due to registration can be observed in Figure 5, the overall reconstruction demonstrates excellent ability to project NFT burden measures into anatomical space, allowing inferences to be drawn about the 3D distribution of NFT pathology in the MTL.

**Figure 5:**
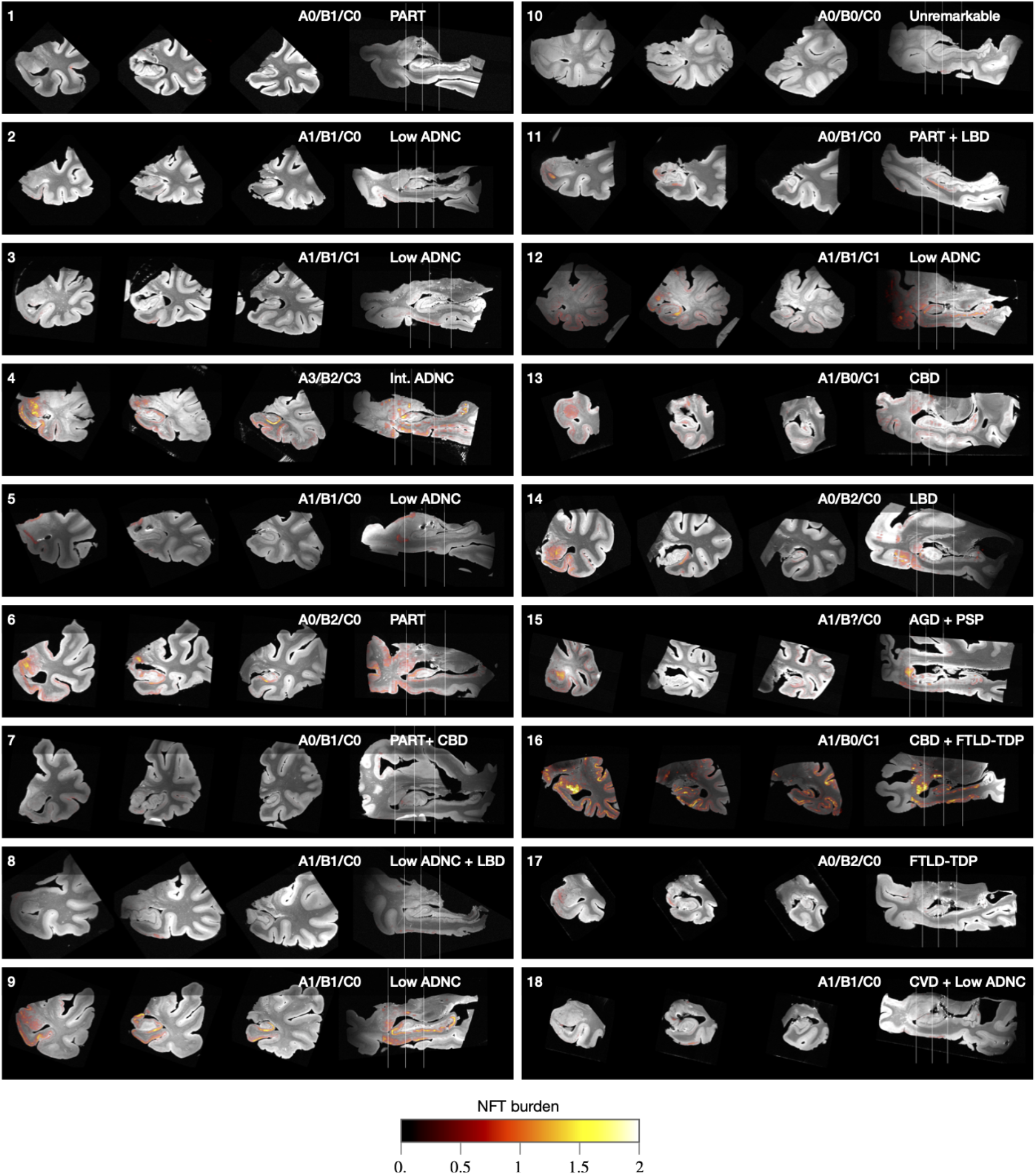
Coronal and sagittal cross-sections of three-dimensional neurofibrillary tangle (NFT) burden maps derived from serial histology and overlaid on the 9.4T MRI scans of medial temporal lobe specimens from 18 brain donors. Each specimen is annotated with the brain donor identifier (cross-referencing Supplemental Table S.1), NIA/AA neuropathological A/B/C staging [36], and primary/secondary pathological diagnoses. The A/B/C staging is from the contralateral hemisphere and encompasses Thal amyloid plaque staging (A0-A3), Braak staging (B0-B3) and CERAD neuritic plaque staging (C0-C3). The coronal slices are through the amygdala, anterior hippocampus and hippocampal body.

In most specimens, the intensity of the NFT burden map agrees with the ordinal Braak (B) tau pathology score obtained in the contralateral hemisphere with the NIA/AA AD staging protocol [36]. One exception is case 16, which had a neuropathological diagnosis of corticobasal degeneration (CBD) with widespread tau pathology and AD Braak stage B0. CBD is a 4R tauopathy and a subtype of frontotemporal lobar degeneration-Tau (FTLD-Tau) [53]. AD Braak staging in FTLD-Tau is challenging due to the difficulty in distinguishing AD neurofibrillary tau pathology from FTLD-associated tauopathy in neurons and glia. Use of AD-specific tau antibody GT38 [32, 33, 61] in this case revealed minimal AD NFTs consistent with a Braak B0 stage of AD tauopathy, while our machine learning approach detected more extensive FTLD related tau. An example IHC slide from case 16 is shown in Supplemental Figure S.12, highlighting the visually similar appearance of these two types of tau inclusions. Future work will examine the ability of our machine learning pipeline to distinguish AD-type NFTs from glial and neuronal tau in FTLD-Tau. In the current paper, we excluded primary FTLD-Tau cases (i.e., cases 13, 15, and 16) from subsequent group-level analysis, since our burden maps in these cases may be capturing non-AD forms of tau.

### 2.4. Group-Level 3D Analysis of NFT Burden Reveals Characteristic Pattern of Distribution in MTL Sub-regions

Using a custom template generation pipeline (Section 5.5.8), donors’ individual 9.4T MRI scans were transformed into the space of a common anatomical template, which itself was matched to an in vivo human brain MRI template (Figure 6, columns 1 and 2). Corresponding NFT burden maps were transformed into template space and the average map of NFT burden was computed using data from 15 non-FTLD-Tau cases ((Figure 6, column 3). This map reveals a clear topographic distribution of NFT burden, with regions of high average burden following anatomical features such as the cornu ammonis and subiculum layers of the hippocampus, the entorhinal cortex, Brodmann area 35, and amygdala. This conspicuous anatomical pattern indicates the success of the various data fusion and registration stages, as consistent failures in any of the stages (IHC to Nissl, Nissl to 9.4T MRI, 9.4T MRI to template) would have resulted in a blurry average map. This average map may offer limited information about the stereotypical progression of NFT burden because of the variability in this sample of brain donors. Figure 6 (columns 4-7) also plots four frequency maps, each indicating how frequently among these 15 cases the NFT burden exceeds a certain threshold, with thresholds corresponding to severe, moderate, mild and rare levels of NFT burden, as inferred from Figure 2(b). Anatomical regions with higher frequency are likely to be involved earlier in the disease.

**Figure 6:**
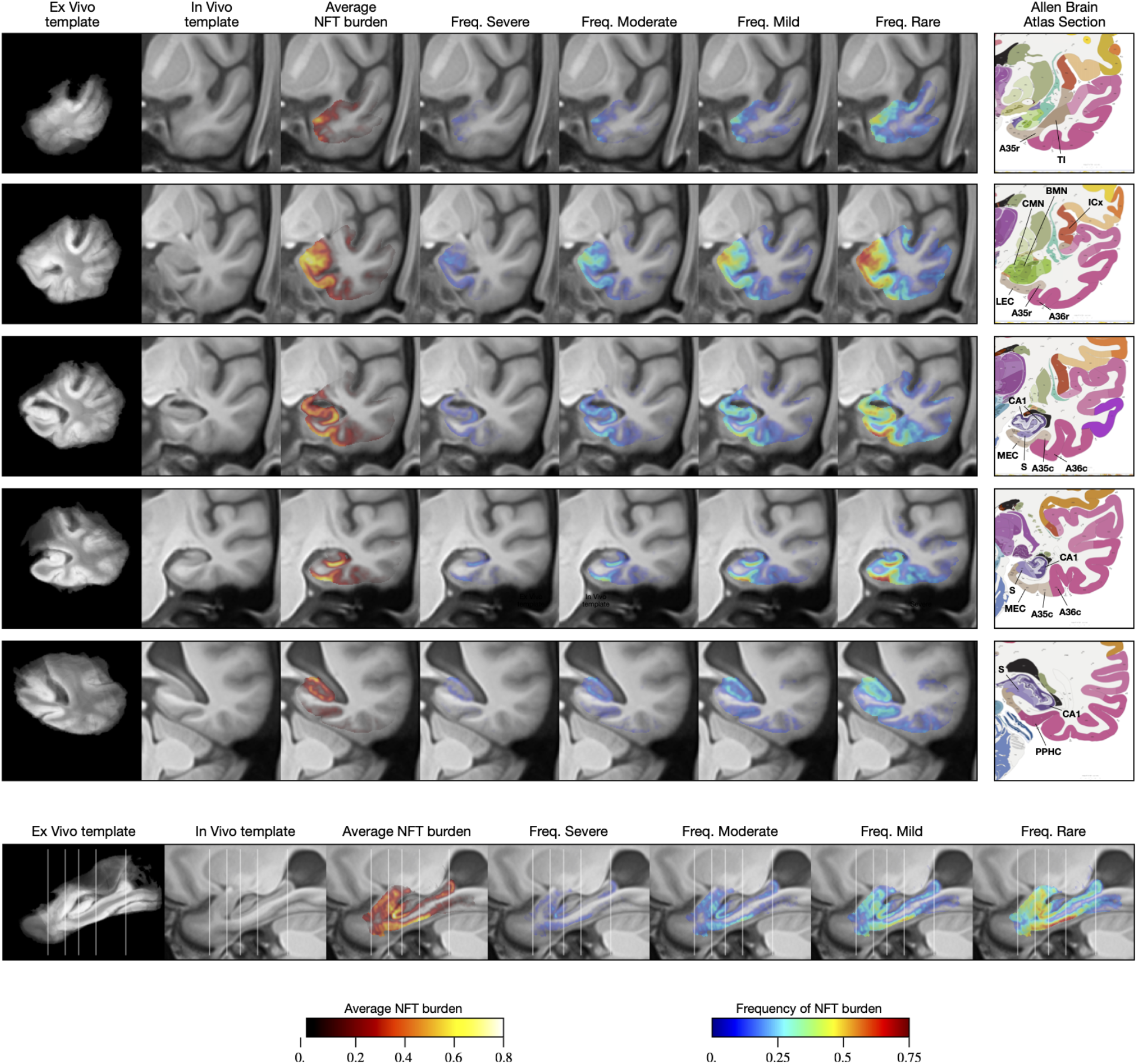
Average and summary maps of tau neurofibrillary tangle (NFT) burden in the space of an anatomical template. These maps use data from 15 of the 18 cases in Figure 5, excluding three cases with primary diagnosis of frontotemporal lobar degeneration - Tau (FTLD-Tau) diagnosis. The top five rows in the figure show coronal cross-sections through the medial temporal lobe at the levels of the temporal pole, amygdala, hippocampal head, hippocampal body and hippocampal tail, respectively. The bottom row shows a sagittal section through the hippocampus. Each row includes a visualization of the average NFT burden map, as well as four frequency maps. The frequency maps at each voxel describe the fraction of cases for which the NFT burden at that voxel was above a given threshold. Thresholds were chosen based on the analysis in Figure 2b and correspond to different levels of pathological burden (>1.0 for “severe”, > 0.5 for “moderate”, > 0.25 for “mild”, >0.1 for “rare”). Coronal sections are also accompanied by roughly corresponding slices from the Allen Brain Atlas [68]. Anatomical abbreviations (following Allen Brain Atlas nomenclature): A35r/A35c: rostral/caudal Brodmann area 35; A36r/A36c: rostral/caudal Brodmann area 36; BMN: mediobasal nucleus of the amygdala; CA1: Cornu ammonis 1 region of the hippocampus; CMN: corticomedial nuclear group of the amygdala; ICx: insular neocortex; LEC/MEC: lateral/medial entorhinal cortex; S: subiculum; PPHC: posterior parahippocampal cortex; TI: temporal agranular insular cortex (area TI).

The sagittal cross-sections of the average burden and frequency maps (bottom row of Figure 6) reveal a clear anterior-posterior gradient, with anterior portion of the parahippocampal gyrus having greater frequency and average burden values than the posterior portion, and similarly greater anterior involvement in the hippocampus. The coronal sections of the maps reveal high frequency of pathology and high average burden levels in Brodmann area 35 (BA35, occupying the medial portion of the perirhinal cortex), entorhinal cortex, amygdala (particularly the medial superior portion, corresponding to the accessory basal and mediobasal nuclei), subiculum and the CA1 subfield of the hippocampus, and the more medial aspect of the temporal polar cortex (possibly corresponding to the very anterior portion of BA35 or to area TI). Cortical regions outside of the entorhinal cortex and BA35 exhibit varying degrees of burden, most prominently BA36 (lateral portion of perirhinal cortex), insular gyri and the inferior temporal gyrus.

Figure 7 plots NFT burden in specific anatomical regions of interest (ROIs) in template space, relative to the burden in BA35. These ROIs were obtained from in vivo MRI atlases [83, 34] matched to the in vivo template. For each ROI defined in the template space, for each of the 15 specimens, we computed the 90th percentile of the NFT burden measure across all voxels in that ROI for which IHC-based measures were available. Figure 7 plots the mean and standard deviation of the ratio of the 90th percentile of NFT burden in each ROI and the 90th percentile of NFT burden in BA35. BA35 was chosen as the reference region since it includes the transentorhinal region (TERC), described by Braak and Braak [15] as the first cortical site of tau pathology in AD. The ROIs with greatest average NFT burden relative to BA35 are the entorhinal cortex (ERC), amygdala, hippocampal subfields anterior subiculum (SUB) and anterior Cornu ammonis field 1 (CA1). The second tier of relatively impacted ROIs includes other areas of the parahippocampal gyrus (BA36, PHC), temporopolar regions (ventral and dorsal area TG), piriform cortex (although due to its proximity to the amygdala, it is likely that this region in the Glasser et al. [34] atlas is picking up NFT burden from the amygdala), and to a lesser extent, insular cortex. Additionally, the ROI analysis highlights the presence of an anterior-posterior gradient in NFT burden, with greater sparing of posterior regions. A more detailed plot of summary ROI NFT burden measures in Supplemental Figure S.13 shows that in individual specimens, including specimens with relatively mild BA35 NFT burden, it is not uncommon for other anatomical regions to exhibit NFT burden similar or greater to that of BA35. Nonetheless, the group analysis quantitatively supports the notion that BA35 and ERC are the regions with greatest NFT burden.

**Figure 7:**
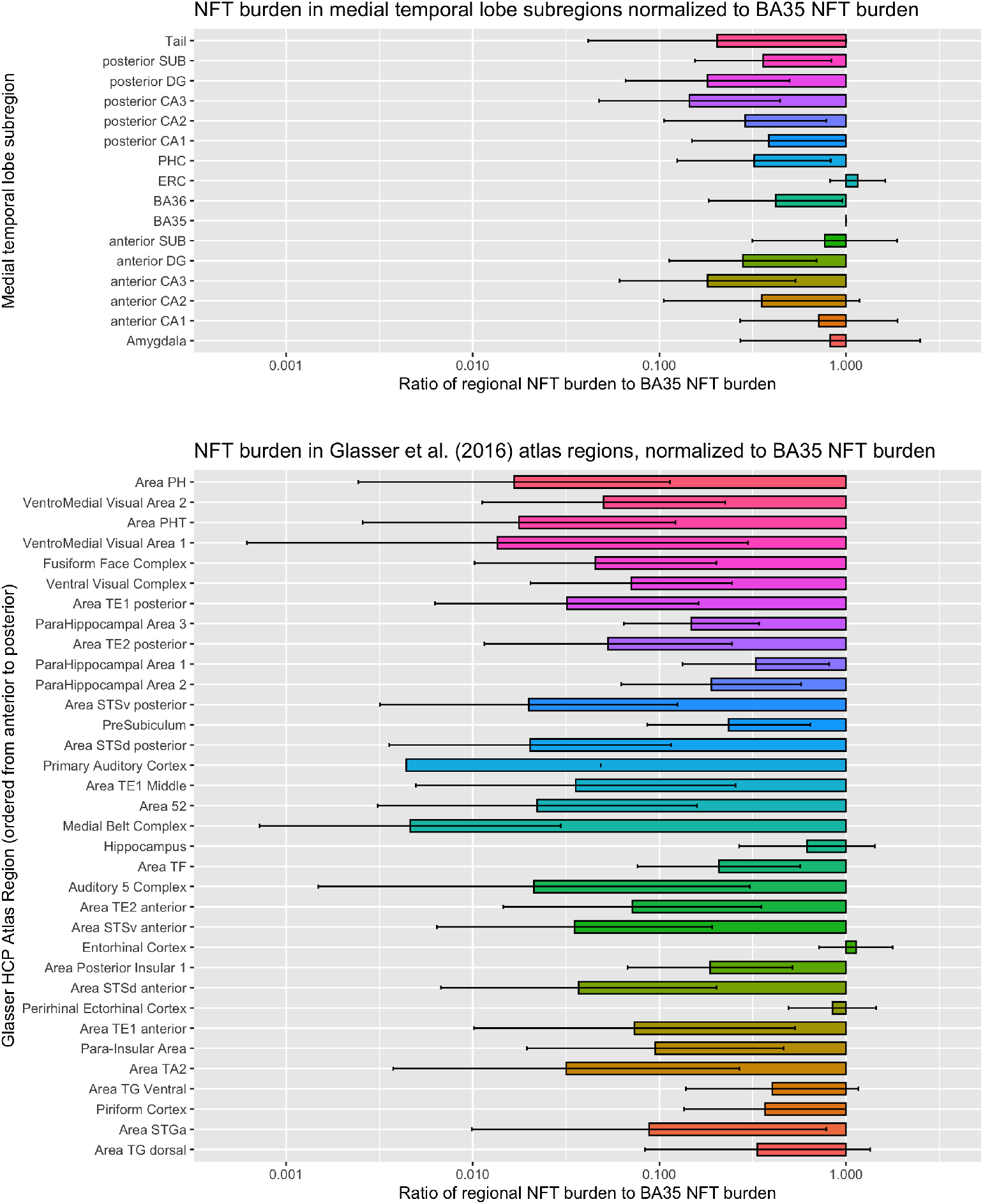
Summary measures of tau neurofibrillary tangle (NFT) burden in two sets of anatomical regions defined in template space. The top plot includes hippocampal subfields and subregions of the parahippocampal gyrus. The bottom plot includes 35 regions from the Glasser et al. [34] Human Connectome Project Multi-Modality Parcellation (MMP) atlas that cover the temporal lobe. For each region, the horizontal bars plot the ratio of NFT burden in that region to the NFT burden in Brodmann Area 35 (BA35), which is thought to encompass the earliest site of cortical NFT formation in Alzheimer’s disease. For each region in each specimen, the summary measure of NFT burden is obtained by taking the 90-th percentile of the regional NFT burden measure. Larger values in the plot indicate greater vulnerability to NFT pathology relative to BA35, while smaller values indicate that regions are spared from NFT pathology relative to BA35. Each set of regions is roughly arranged in the order from more posterior (top of plot) to more anterior (bottom of plot), indicating a presence of a posterior-anterior gradient in relative NFT burden, with posterior structures more spared relative to the anterior structures. These plots use data from 15 of the 18 cases in Figure 5, excluding three cases with primary FTLD-Tau diagnosis.

## 3. Discussion

### 3.1. 3D Maps of Tau NFT Pathology Generated for 18 Brain Donors

To our knowledge, this work is the first to perform 3D mapping of tau NFT pathology in a sizable collection of human brain specimens (*n* = 18). In a recent preprint, Alegro et al. [4] describe a pioneering pipeline for 3D tau density maps generation in whole brains and present results for two brain specimens. They use deep learning to extract and quantify tau inclusions from anti-tau immunohistochemistry slides and use registration to ex vivo MRI to reconstruct 3D maps. Alegro et al. [4] perform ex vivo MRI and CT *in situ*, an advantage over the current work since this approach eliminates MRI artifacts due to air bubbles and makes co-registration with *in vivo* MRI simpler. However, *in situ* scanning is not always feasible to integrate into existing autopsy protocols, including at the brain banks used in our study. Like the current paper, Alegro et al. [4] also utilize 3D printing, but for a different purpose, to create customized containers to minimize brain distortion during formalin fixation. While the whole-brain mapping and *in situ* imaging are impressive advantages of the [4] pipeline, the amount of manual interaction in their pipeline is greater, requiring manual initialization for spline-based deformable registration for each histology slide. Tward et al. [72] describe a new deformable registration approach for aligning serial anti-tau immunohistochemistry slides to ex vivo MRI without a need for intensity transfer, and implement it for several tissue blocks from one brain donor. Tward et al. [72] also use deep learning to generate tangle heat maps; however they use a simpler patch classification strategy (assigning each patch a tangle/non-tangle label, as opposed to weakly supervised learning used in this paper) and do not evaluate resulting heat maps against manual tangle counts or semi-quantitative scores. Given that previous tau 3D mapping efforts [72, 4] were applied to only one or two specimens, the current work is the first to analyze histologically-derived 3D maps of tau pathology in a group template space and derive average and summary maps of NFT pathology burden (Fig. 6). The fact that we were successful in applying our approach to 18 specimens also speaks to the generalizability and scalability of our approach to other individuals with other neurological conditions.

### 3.2. 3D Pattern of Tau NFT Burden Recapitulates Earlier Neuropathological Studies of NFT Spread in the MTL

The patterns of tau NFT distribution in individual specimens (Figure 5) and in the group template space (Figure 6) are consistent with earlier histological literature that characterized NFT spread in the human brain. For instance, Arnold et al. [5], quantified NFT density in 39 cortical regions in patients with clinically intractable dementia confirmed pathologically as AD. Greatest average NFT density was found in BA28 (ventral entorhinal cortex), subiculum/CA1, BA38 (temporopolar cortex), BA35, posterior parahippocampal areas TF and TH, amygdala, and BA51 (piriform cortex). Essentially the same areas come out at the top of our analysis. However, whereas in Arnold et al. [5], the NFT density in many regions was only slightly lower than in the most affected regions (e.g., 2.8 in the anterior insula vs. 3.9 in BA28), in our analysis the distribution is less uniform, with most regions outside of the medial temporal lobe and temporopolar cortex having NFT burden reduced threefold or more relative to BA35. These similarities and differences are consistent with the brain donors in our study being in earlier stages of disease progression than the donor cohort in [5].

Arguably, the most influential work on characterizing the spread of NFT pathology is by Braak and Braak [16, 17], whose six-stage staging system is used universally for neuropathological diagnosis of AD [36]. Braak and Braak [16] used the term “transentorhinal” to describe the early stages I and II, and argued that the earliest manifestations of NFT pathology are largely confined to the transentorhinal region (medial BA35), with some involvement of the entorhinal cortex as well. While many of the brain donors in our cohort were classified as B1 (corresponding to the “transentorhinal” stages I and II), our 3D analysis suggests a more widespread distribution of NFT burden. In a number of cases with mild transentorhinal involvement, the amygdala, anterior CA1 and subiculum, and anterior temporal pole regions exhibit NFT burden on par or even greater than in BA35 (Supplemental Figure S.13, Figures 5, 6). This discrepancy could be attributed to asymmetry (Braak staging was performed using the contralateral hemisphere) or limited sampling of tissue for Braak staging. Overall, however, our results suggest that in studies of early AD, a broader focus beyond the transentorhinal region, and inclusive of hippocampal subregions, amygdala, and temporopolar cortex may be warranted.

The vulnerability of temporopolar cortex and amygdala to NFT pathology has been described in the neuropathology literature [47, 6, 30]. Kromer Vogt et al. [47] identified the basomedial and cortical nuclei of the amygdala as having highest NFT density. Interestingly these regions are strongly connected to regions typically thought to harbor early NFT pathology, such as the ERC and PRC [57]. Our group-level 3D map appears to reveal elevated frequency of supra-threshold NFT burden in the cortical and basomedial nuclei relative to the rest of the amygdala (Figure 6), although the precise distribution of tau burden in the amygdala is difficult to infer in the groupwise template, and will require a focused follow-up study with cytoarchitecture-based labeling of amygdala sub-nuclei. In the temporopolar cortex, Ding et al. [30] reported greatest density of anti-tau labeled neurons in the anterior BA35, followed by areas TG and TI, and to a lesser extent anterior BA36. The most anterior slice in Figure 6 reveals two peaks in the NFT frequency maps whose locations are consistent with anterior BA35 and area TG as defined by Ding et al. [30]. This finding suggests that the early tau pathology of BA35, as well as the lateral entorhinal area, which also overlapped high levels of NFT in the frequency maps, extend much more anterior in these regions than usually considered in segmentation protocols measuring these areas on in vivo studies of AD [83].

### 3.3. *3D Pattern of Tau NFT Burden Exhibits Similarities with* In Vivo *Neuroimaging*

Examining the NFT burden maps in Figures 5 and 6 along the sagittal axis reveals an anatomical pattern that cannot be visualized with conventional 2D histological analysis. Indeed, the regional analysis of prior histological studies describes NFT involvement in monolithic terms whereas the current 3D mapping suggests gradients of involvement even within regions. Namely, we observed a directional gradient in the NFT burden, with greater involvement of anterior areas of the parahippocampal gyrus (anterior hippocampus, entorhinal cortex, BA35, amygdala). Ranganath and Ritchey [58] hypothesized that memory function is subserved by two MTL networks, anterior temporal (AT) and posterior medial (PM) networks. The AT network, which involves the perirhinal cortex (BA35), anterior hippocampus, lateral entorhinal cortex, amygdala, ventral temporopolar cortex, and orbitofrontal cortex, is linked to familiarity-based recognition memory, social cognition, and semantic memory; while the PM network involves medial entorhinal cortex, posterior hippocampus, parahippocampal cortex, and retrosplenial cortex, and is linked to episodic, or recollection-based memory, spatial navigation, and situational models or schemas [58]. It was proposed in this model, that semantic dementia best maps on to the AT system while AD most prominently involves the PM network, which overlaps with the default mode network, but not completely to the exclusion of the AT. However, the 3D distribution of NFT burden in our results, with strong involvement of anterior structures, aligns closely with the AT network, with BA35, amygdala, anterior CA1/subiculum, and temporopolar regions, particularly the anterior extension of BA35, having most pronounced NFT burden. In fact, a number of studies have suggested alterations in cognition in very early AD more linked to the AT network, including object short and long-term memory, familiarity, and semantic memory [44, 76, 27, 52, 12]. One explanation for the relative sparing of the PM system in our 3D NFT data may be that non-tangle forms of tau pathology, including neuropil threads and tau in neuritic plaques, contribute to the deterioration of the PM network in AD. Another explanation may be that in the presence of extensive beta-amyloid pathology, the distribution of NFTs is more posterior than in the majority low beta-amyloid cases that compose our cohort, although 2D histology studies have found PART to have a very similar distribution to that of AD [25]. Future analysis including more cases with more pronounced beta-amyloid pathology and examining additional tau species may shed light on AT and PM network involvement in AD and PART.

That said, *in vivo* studies of the AT and PM systems utilizing tau and amyloid PET are largely consistent with our 3D map of NFT burden. Schwarz et al. [64] identify patterns of ^18^*F*-AV-1451 tau tracer uptake associated with progressive Braak stages, and report the pattern in the “transentorhinal” stages I and II impacting the transentorhinal region (which, given the resolution of PET encompasses ERC and BA35), hippocampus, and amygdala. Lowe et al. [50] compared ^18^*F*-AV-1451 uptake across brain regions in cognitively impaired and unimpaired individuals with and without cognitive decline, and report amygdala and temporal pole as the two regions with greatest tau uptake in cognitively unimpaired, amyloid-negative group, consistent with the high NFT burden in these regions in our analysis. They also report findings of elevated tau uptake in regions outside of the MTL in unimpaired, amyloid-negative individuals, which is consistent with our findings of elevated NFT burden in a number of cortical regions even when NFT burden in BA35 was low. Maass et al. [52] examined ^18^*F*-FTP-tau PET uptake in AT and PM network regions in normal aging and AD, and found elevated tau PET uptake in AT regions relative to PM regions in both aging and AD, whereas amyloid PET uptake was greater in the PM network regions and concluded that “posterior-medial regions are affected by tau later in the course of the disease and do not ‘catch up’ while anterior-temporal tau accumulation accelerates further”. This finding of increased AT vs PM tau is consistent with the anterior-posterior gradient of NFT burden in our ex vivo analysis. It is worth noting that given the greater amyloid burden in the PM network, it is possible that the tau in these regions are disproportionately neuritic while the AT network may reflect NFTs, as measured here. Given these general similarities between patterns of tau distribution in our NFT burden maps and in vivo tau PET uptake patterns, it would be promising in future research to carry out analyses that directly contrast these data on a voxel-wise basis.

### 3.4. A Largely Automated Pipeline for 3D Mapping of Proteinopathies from Serial Histology and MRI

Our approach incorporates a complex image analysis pipeline that, in contrast to our earlier work on MRI/histology co-registration [1], is largely automated, with only a few manual initialization steps (i.e., initial alignment) that involve whole specimens or whole blocks, as opposed to individual 2D slides. As a result, the human and computational effort to create a new reconstruction for a single specimen is significantly smaller (^~^1 day alternating between manual steps and automated processing) than the time to perform imaging and histology procedures. Among the novel aspects of the pipeline is the use of custom 3D printed molds to guide tissue cutting and reduce the number of degrees of freedom for histology-to-MRI image registration. Mold-guided tissue cutting also ensures parallel cuts of the tissue and better sectioning, with less tissue lost at the ends of each block. This concept was previously proposed and implemented for marmoset and rat brains [51, 14], but to our knowledge this is the first application custom 3D printed molds to guide registration in a significant number of human brain specimens.

Our pipeline uses *intensity transfer* to account for differences in appearance between histology and MRI during image registration (Supplemental Figures S.6, S.7, Section 5.5.3). Intensity transfer allows registration to take advantage of existing robust image registration tools, rather than requiring customized image similarity metrics as in other recent work on MRI/histology co-registration [56, 37, 72]. While intensity transfer using generative adversarial networks and other deep learning techniques has been applied widely in medical imaging [80], its use in histology-MRI registration is limited, likely due to large differences in image resolution. Ours is the first such approach to leverage unsupervised deep clustering [22] to generate a rich set of lower-resolution features from whole-slide histology images, which can then be mapped to MRI-like appearance using robust least square fitting.

Another feature of our pipeline is to use weakly supervised learning with modified WildCat algorithm [31] to infer regional pathology burden. This approach yields more detailed maps than pure image classification, in which a CNN is used to assign each image patch a single vector of object membership probabilities and whole-slide maps are generated by a sliding window approach (such a strategy was implemented by Tang et al. [69] for quantification of amyloid plaque and cerebral amyloid angiopathy burden). Yet it is much less onerous to train than full-blown segmentation of individual tangles [65, 78, 4], which requires tangles and other objects to be segmented in the training data, and may be unnecessary when the objective is to derive maps of regional NFT burden at the resolution of the ex vivo MRI. In unpublished data, rapid generation of training data allowed us to easily extend this approach to quantifying the burden of Lewy bodies and TDP-43 intercellular inclusions on archival whole histology slides.

### 3.5. Limitations and Future Work

A strength of the presented pipeline is that it allows 3D NFT density maps to be computed in MRI space with relatively little training effort (drawing boxes) and relatively little manual input for registration (seeding affine registrations). The accuracy of the registration is on the order of 1-2 ex vivo MRI voxels on average in the four anatomical regions examined, and visually, it appears that histology and MRI are well aligned through most of the temporal lobe. While this error is significant on the scale of individual tangles and even cortical layers, it is quite small relative to the resolution of in vivo MRI or PET and to errors encountered for in vivo MRI registration, e.g., between individual MRI scans and population templates. Hence, for applications where 3D patterns of NFT burden are needed to inform in vivo MRI analysis (e.g., to derive hot spots in which to measure atrophy linked to tau pathology), the histology-to-MRI registration errors can be considered almost negligible.

Even though the Wildcat classifier achieves >95% accuracy for individual tangle/non-tangle patches, at the whole-slide level, WildCat classification is not devoid of error. In particular, our training set includes a very small number (<2%) of astrocytic tau inclusions, which can be similar in visual appearance to tangles. Indeed some areas of age-related tau astrogliopathy (ARTAG) [45] are incorrectly identified as areas of high NFT burden by the algorithm. Overcoming this limitation requires extending the training set with a large number of ARTAG examples and ensuring that the high accuracy of WildCat is maintained. ARTAG misclassified as NFT may contribute to elevated measures of NFT burden in some cortical regions and amygdala, where ARTAG is more common [45]. Our algorithm also does not distinguish tangles from pretangles, which may lead to confounding of different stages of NFT pathology. However, visual assessment of the data suggests that ARTAG and pre-tangles are only present in a small fraction of cases and location is not consistent from case to case, making it likely that their impact on NFT burden maps reported here is relatively minor. Lastly, the WildCat algorithm was not trained to differentiate NFTs from similarly appearing 4R-tau inclusions in FTLD-Tau (Supplemental Figure S.12), necessitating the exclusion of FTLD-Tau cases from group analysis.

The groupwise registration used a conventional population-based template approach [42, 9], whereas in earlier work we showed that a method initialized with shape-based registration of semi-automatic segmentations of anatomical regions of interest (hippocampus, MTL cortex) results in better groupwise registration of these regions [1, 59]. However, given the large anatomical extent considered in this study (entire temporal lobe, not just MTL) and variation in extent between individuals, this shape-based approach would be difficult to implement. Some improvements to registration can likely be realized by incorporating more specimens and considering more advanced groupwise registration strategies [81, 3]. A related limitation is that the anatomical labels for the analysis were derived from in vivo atlases, and may not match individual anatomical structures well in group template space. We are currently generating cytoarchitecture-guided segmentations of MTL subregions in the specimens in this study, and intend to conduct more granular analyses, including analyses linking NFT burden to regional alterations in MTL cortical thickness in future work. Thus, the present work serves as a foundation upon which future work can build to characterize the relationships between tau and other proteinopathies in a variety of forms (e.g. ARTAG versus NFTs) and structural changes in the MTL. Such future studies will enhance our understanding of the local neurodegenerative consequences of these pathologies.

## 4. Conclusions

We presented a technique for combining data from ex vivo MRI and serial histology to generate 3D maps of NFT density in MRI space. These maps help provide information about the distribution of NFT pathology in the region of the cortex where it first emerges in AD, and may help inform the field about the patterns of NFT pathology spread, as well as support development and validation of better AD biomarkers.

## Acknowledgments

We gratefully acknowledge the generosity and altruism of the brain donors and their families. We thank the members of the brain banks at the University of Pennsylvania Center for Neurodegenerative Disease Research and at the University of Castilla La Mancha at Albacete for their contributions to this research. This work was supported by NIH grants P30 AG010124, R01 AG056014, and R01 EB017255.

## Author Contributions

Conceptualization: P.A.Y., L.E.M.W., D.A.W., D.J.I., R. I.; Methodology: P.A.Y., S.R. S.P., J.W., L.Y.H., J.L., N.V., L.X., M.D.T., G.M., S.R.D., R.I.; Software: P.A.Y., S.R., J.W., J.L., N.V; Validation: P.A.Y., S.L.; Investigation: P.A.Y., M.M.I.d.O.M., R. I., S.L., S.R., M.L.B., S.P., W.L., M.D.T., K.P., G.M., D.T.O., L.E.M.W., D.J.I., R.I., M.M.A-J, E.A-P, P.M.R, M.M.L., C. D-R.P; Resources: J.L.R., T.S., M.G., E.B.L., J.Q.T; Data Curation: P.A.Y., S.R., M.L.B., J.L., N.V., L.X., M.D., S.C., L.M., R.d.F., S.R.D., D.T.O., D.J.I., R.I., M.M.A-J, E.A-P, P.M.R, M.M.L., C. D-R.P., S.C.S., J.G.D.G., M.C.P., FJ. M.R.; Writing – Original Draft: P.A.Y.; Writing – Review & Editing: P.A.Y., M.M.L., S.R., L.X., S.R.D., E.B.L., J.Q.T., D.T.O., L.E.M.W., D.A.W., D.J.I.; R.I.; Supervision: P.A.Y., R.I.; Funding Acquisition: P.A.Y., D.A.W., J.Q.T.

## 5. Online Methods

### 5.1. Data and Code Availability

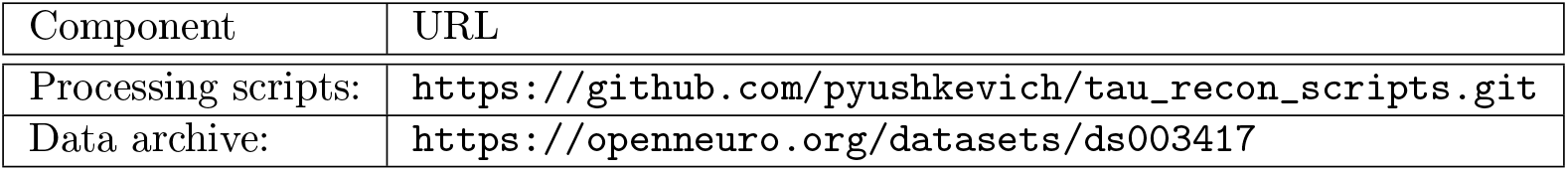

### 5.2. Specimen Preparation and Imaging

Human tissue specimens acquired in the cohort of 18 brain donors summarized in Table 1 and Supplemental Table S.1 underwent a sequence of preparation and imaging procedures summarized in Fig. 1. Each step is described below.

#### 5.2.1. Hemisphere Selection and Fixation

For each brain donation, tissue from one brain hemisphere was used for imaging (MRI and histology) and the opposite hemisphere was sampled for diagnostic pathology following the NIA-AA protocol [36]. At UCLM, left vs. right hemisphere was selected at random, while at UPenn, hemispheres selected for imaging were those for which contrast in the hippocampal region was best on antemortem in vivo MRI (if available), and at random otherwise. Different fixation procedures are followed by the UCLM and UPenn brain banks. UCLM performs fixation in situ before brain removal at autopsy by pumping 4% paraformaldehyde in 0.1M phosphate buffer through the carotid artery. UPenn performs fixation in the imaged hemisphere after the brain is removed. In both cases, hemispheres stay in fixative for 4+ weeks.

#### 5.2.2. MTL Specimen Preparation for MRI

After fixation, a specimen containing the intact MTL is dissected from the hemisphere. The criteria for dissecting the MTL specimen are that it must fit into a cylindrical holder with 50mm inner diameter and that the MTL structures, including amygdala, temporal pole, perirhinal and parahippocampal cortex, and the tail of the hippocampus are intact. An example of the excised region is shown in Figure S.4. The surface of the MTL specimen is stripped of pia mater. The specimen is placed in a desiccator for several hours to remove surface fluid and then placed in the specimen holder filled with Fomblin, a MRI-neutral recycled industrial oil. To reduce air bubbles that cause MRI artifact, the specimen is placed in the holder and the holder lid is closed while submerged in a Fomblin bath. The holder is then agitated overnight to help extract remaining air bubbles from the sulci.

#### 5.2.3. 9.4 Tesla and 7 Tesla MRI

All MRI procedures were performed at UPenn. Each intact MTL specimen was scanned overnight on a 9.4T Varian animal MRI scanner with a horizontal 300mm bore, 70mm inner diameter quadrature transmit/receive volume coil, and Bruker console. The scans use the product 2D multi-slice spin echo sequence (TR=9s, TE=23ms, 48×128mm field of view, 240 axial slices, 27 averages, 16 hour scan time, with some deviations in the protocol for individual specimens) resulting in an 0.2 x 0.2 x 0.2 mm^3^ native resolution image with a mixture of T2 and proton density weighting. In all except for one scan, the entire MTL specimen was included in the field of view.

While the 9.4T scanner offers high signal to noise ratio for imaging MTL specimens, as previously reported [1], it suffers from geometric distortions due to the non-linearity of the magnetic gradient field that increases towards the ends of the sample. In order to 3D print cutting molds that fit the specimens closely, each excised MTL specimen was additionally scanned using a 7 Tesla Siemens Terra MRI scanner (Siemens Inc, Erlangen, Germany) in a custom 70mm inner diameter quadrature transmit/receive volume coil using the product 3D T2-weighted sequence (T2-SPACE, TR=3000ms, TE=380ms, 4 averages, 19 min scan time) with 0.4 x 0.4 x 0.4 mm^3^ resolution. The vendor’s on-scanner correction for gradient-field non-linearity was applied to the 3D volumes, and a grid phantom was used to confirm low geometric distortion on the 7T scanner (<1mm). Examples of 7T and 9.4T scans are shown in Fig. 1.

#### 5.2.4. 3D Printed Cutting Mold

To achieve maximal congruence between MRI and histological sectioning, a custom cutting mold (Fig. S.4) is 3D printed for each specimen using a model created semi-automatically from the 7T MRI scan (as detailed in Supplemental Section Appendix A.1). The mold tightly fits the MTL specimen and has a series of parallel slits at 4mm intervals that run roughly orthogonal to the main axis of the hippocampus. By using these slits to section the tissue into blocks, we ensure that the plane of histological sectioning relative to the 7T MRI scan is known, thus reducing the complexity of subsequent registration between MRI and histology. Additionally, compared to conventional tissue blocking, the cutting mold ensures that the blocks are cut with parallel sides, which, we observed, greatly reduces the amount of tissue discarded at the ends of the blocks during histological sectioning.

#### 5.2.5. Serial Histology and Immunohistochemistry (IHC)

All histology procedures were performed at UCLM, with the exception of digital scanning, which was performed at multiple sites. Using the custom mold, specimens were cut into 20 mm thick blocks, with most specimens yielding 4 blocks. Cryoprotected blocks were frozen using dry ice and sectioned using a sliding microtome coupled to a freezing unit into 50 *μ*m sections, with no gaps between sections. Before cutting each section, a digital photograph of the block were taken using a mounted overhead camera (called *blockface images*).

Each section, and its corresponding “blockface” image, were individually and spatially identified. Every tenth section (sections 10, 20, 30,…) was stained for the Nissl series using the thionin stain. Every twentieth section (19,39,59,…) was stained using AT8, a human phosphorylated tau antibody IHC stain, and were counterstained for Nissl. Thus, Nissl stained sections were at 0.5mm intervals (^~^40 per block) and anti-tau sections were adjacent to the Nissl sections and at 1mm intervals (^~^20 per block). Additional sections (36, 37, 38, 76, 77, 78,…) were designated for additional IHC stains (α-synuclein, TDP-43, β-amyloid). However, these additional stains were only obtained when there was evidence of the respective pathologies in the contralateral hemisphere, which underwent detailed neuropathological evaluation with staining of multiple regions for α-synuclein, TDP-43, β-amyloid, and tau based on the NIA/AA ADNPC criteria [36]. These additional IHC stains are not studied in the current paper.

Sections were mounted on 75 mm × 50 mm glass slides and digitally scanned at 20X resolution (Fig. 1). For three specimens, scanning was performed on the Aperio ScanScope digital scanning microscope; for the rest, a Huron LT120 digital scanning microscope was used. Scans were uploaded to an in-house created cloud-based digital histology archive that supports web-based visualization, anatomical labeling, and machine learning classifier training.

### 5.3. Derivation of NFT Burden Maps using Weakly Supervised Learning

#### 5.3.1. Training Data Generation

Each of the 12 raters participating in the Tanglethon events was assigned a set of AT8-stained anti-tau IHC slides and asked to label a large number of “tangle-like” objects (tangles and pre-tangles) and “not tangle-like” objects (tau neuropil threads, astroglial tau, tau coils in the white matter, normal tissue, slide background, artifacts, tissue folds). Each object was labeled by placing a box around it in the web-based system and assigning a categorical label. Raters were trained and supervised by two experts in neurodegenerative histopathology (DJI, DTO). The experts reviewed objects that raters were unsure how to label, and objects that elicited uncertainly among the experts were assigned the “ambiguous/uncertain” label. Using this expert-supervised crowdsourcing approach, over 11,000 patches were extracted and labeled from 176 slides in 6 MTL specimens. To generate training, validation and testing data for WSL, patches of size 512 × 512 pixels centered on each object were extracted from the full-resolution IHC slides and grouped into tangle-like (*n* = 4975) and non-tangle-like classes (*n* = 6386), with ambiguous/uncertain patches excluded. Examples of patches in each class are shown in Supplemental Figure S.2.

#### 5.3.2. Derivation of NFT Burden Maps using the WildCat Algorithm

WildCat [31] is a weakly supervised learning algorithm formulated as a convolutional neural network (CNN). It is derived from the ResNet image classification CNN [35], but replaces the last fully connected layer of ResNet with a class pooling layer and a spatial pooling layer. These layers are 1/16 of the size of dimension of the input image and provide a low resolution heat map that was shown in [31] to localize real-world objects in scenes. For example, in a photo containing two cats, the trained Wildcat classifier will generate heat maps in the class pooling layer with peaks that correspond to the cats’ locations. In [31], these heat maps are post-processed using a conditional random field (CRF) [48] to perform pixel-level segmentation. To reduce computational cost, and because CRF may have trouble differentiating tangles from similarly-appearing non-tangle tau inclusions (e.g., tau threads, tufted astrocytes), we instead modified WildCat with a U-net like structure, adding upsampling layers with skip connections [62] to ResNet layers to generate higher-resolution class pooling and spatial pooling layers. The result is a heat map that is 1/2 of the input image size, as opposed to 1/16 in the original WildCat formulation. Sample patches and corresponding heat maps are shown in Supplemental Figure S.2, showing that the heat maps colocate with tangles, while having a more blurry appearance.

#### 5.3.3. Evaluation of NFT Classification Accuracy

WildCat is an image classification CNN modified to generate spatial heat maps that indicate which regions of the input image contribute to the classification decision. To evaluate the classification accuracy of WildCat for tangle detection in specimens unseen during training, we performed leave-one-out crossvalidation experiments using anti-tau IHC slides from the six specimens annotated during the Tanglethon. In each experiment, we trained a WildCat model using 3000 randomly selected patches from five specimens (2000 for training, 1000 for validation) and evaluated the patch-level accuracy of the trained model using 1000 randomly selected patches from the remaining specimen. WildCat pooling parameters were set to *k^−^* =0, *k^+^* = 0.02, *α* = 0.07, *M* = 4 (see [31]). The models were trained over 30 epochs, and the model parameters from the epoch with the best validation set classification accuracy were used subsequently.

#### 5.3.4. Generation of Patch-Level and Whole-Slide NFT Burden Maps

To obtain an NFT burden heat map for an IHC image patch, we apply the trained WildCat model to the patch, extract activation maps from the class pooling layer, subtract the activation map for the non-tangle-like class from the activation map for the tangle-like class, and threshold the difference at zero. Examples of difference maps for sample tangle-like and non-tangle patches are shown in Supplemental Figure S.2.

This provided a “NFT burden heat map” shown in Fig. 2a.

We found WildCat output to be consistent across input patches of different size.

#### 5.3.5. Validation of NFT Burden Maps vs. Conventional Measures

The 48 boxes used to validate WildCat against manual NFT counting (Section 2.1.3) and semiquantitative ordinal ratings (Section 2.1.3) were extracted from anti-tau ICH as follows. Author S.L. traced multiple curves through the gray matter in 12 slides in 3 specimens in the web-based histology annotation system. In each slide, 50 boxes of 2048 x 2048 pixels overlapping these curves were extracted at random. Author D.J.I. identified 4 boxes per slide that were representative of the range of pathology in that slide,. Author R. It. outlined individual tangles in each of the selected boxes, which were reviewed by D.J.I. D.J.I. also assigned a subjective ordinal rating of tau pathology severity to each box using a standardized scale [36] with categories “0 (None)”, “0/1 (Rare)”, “1 (Mild)”, “2 (Moderate)”, and “3 (Severe)”.

A single WildCat model was trained on all 6 specimens included in the Tanglethon, using the same parameters as in the above cross-validation experiments. The model was applied to generate NFT burden heat maps in the 140 boxes. From the class pooling layer, we subtracted the activation map for the nontangle-like class from the activation map for the tangle-like class and thresholded the difference at zero. This provided a “NFT burden heat map” shown in Fig. 2a. This heat map was integrated over each box, providing a quantitative measure of NFT burden in the box. Summary NFT burden was compared with manual tangle counts using non-parametric correlation tests (Spearman *ρ*, Kendall *τ*). The ability of summary NFT burden measures to discriminate between subjective ordinal rating categories was evaluated using the Mann-Whitney U-test.

#### 5.3.6. Generation of Whole-Slide NFT Burden Maps

Whole-slide NFT burden maps were obtained by breaking the slide up into non-overlapping windows of 4096 x 4096 pixels, running WildCat trained on the full set of 6 specimens on these windows, subtracting the non-NFT class activation map from the NFT class activation map, thresholding at zero, and additionally downsampling by the factor of 4, resulting in a 512 x 512 pixel heat map. These heat maps were tiled to generate whole-slide heat maps. An example of a whole-slide NFT burden map is shown in Supplemental Figure S.3. These maps were then transformed into MRI space using a multi-stage 3D reconstruction pipeline described in the next Section.

### 5.4. 3D Reconstruction Pipeline

### 5.5. Image Analysis Pipeline Overview

Transforming NFT burden heat maps into MRI space is a complex multi-stage process, with the main stages summarized in Table 2 and detailed in the sections below. A more detailed flowchart of the processing and registration steps undergone by the five imaging modalities collected in this study (7T MRI, 9.4T MRI, blockface images, Nissl histology, anti-tau IHC) is shown in Supplemental Figure S.4. A common feature in the pipeline is that when dealing with multi-modality registration problems (between MRI and blockface images or between MRI and Nissl histology), we transform the intensity of the blockface or histology images to have an “MRI-like” appearance. This transformation allows us to perform image deformable registration using the normalized cross-correlation (NCC) image similarity metric [8], which we found to result in much better matching than the mutual information metric, which is formulated for multi-modality image registration.

**Table 2:** Summary of the steps in the MRI/histology reconstruction pipeline. Each stage of the pipeline is described in the text.

| Pipeline Stage | Section | Scope | Purpose |
| --- | --- | --- | --- |
| MRI Processing | 5.5.1 | whole specimen, 3D | Correct for geometric distortion in 9.4T scans via deformable registration to the 7T MRI. Includes registration box **A** in Fig. S.4. |
| Alignment of MRI to 3D reconstructed blockface volume | 5.5.2 | block, 3D | Bring MRI and histology stacks into approximate alignment leveraging geometric information from 3D printing models. Corresponds to box **B** in Fig. S.4. |
| Nissl intensity remapping | 5.5.3 | slide, 2D | Reduce the resolution of Nissl images to be on the same order of magnitude as MRI resolution and generate appearance that is comparable to that of MRI, allowing registration between MRI and Nissl slides using conventional image similarity metrics. |
| Nissl to MRI registration | 5.5.4 | block, 2D/3D | Align and warp each individual slide to the corresponding cross-section of the MRI volume, correcting for distortions during histological processing and generating a 3D histology volume. Corresponds to boxes **D**, **E** in Fig. S.4. |
| IHC to Nissl registration | 5.5.5 | slide, 2D | Map corresponding locations from tau IHC slides to adjacent Nissl slides using piecewise diffeomorphic registration. Corresponds to box **C** in Fig. S.4. |
| Final 3D reconstruction | 5.5.7 | whole specimen, 3D | Map NISSL and IHC images and derived NFT burden maps into the whole-specimen MRI space. |
| Multi-subject registration | 5.5.8 | all specimens, 3D | Combine NFT load maps from multiple specimens in a common anatomical frame to allow group-level comparisons. |

#### 5.5.1. MRI Processing

MRI scans underwent intensity inhomogeneity correction using the N4 tool [71]. Each high-resolution 9.4T scan was co-registered to the corresponding 7T scan using affine registration (manually initialized using ITK-SNAP) and deformable registration with extensive regularization of the transformation field (registration details in Supplemental Table S.4A). The deformable registration accounts for the non-linear geometric distortion caused by gradient field inhomogeneity on the 9.4T scanner, and increased regularization is used because of known smooth nature of these distortions. Visual inspection confirmed excellent alignment between 7T and 9.4T scans, except in areas where semi-detached pieces of tissue moved between scans or in areas of MRI artifact. In the remainder of this section, we will refer to the 9.4T MRI scan that has been warped into the space of 7T MRI and resampled to 0.2 × 0.2 × 0.2 mm^3^ resolution as the “distortion-corrected 9.4T MRI” (see Fig. S.5).

#### 5.5.2. Alignment of MRI to the 3D Reconstruction of Blockface Images

##### Blockface Reconstruction

Blockface images were cropped, removing excess background, and downsampled to approximately match the resolution of the 9.4T MRI. Blockface images were stacked in 3D with the interslice spacing set to 0.05mm. We observed that, occasionally, the camera had shifted slightly when taking blockface photographs. Such shifts were automatically detected and corrected via rigid registration when forming 3D stacks. An example blockface image stack is shown in Supplemental Figure S.5a.

##### Blockface Intensity Remapping

To facilitate registration with the MRI, blockface images were transformed to an “MRI-like” appearance, i.e., black background, dark white matter, and lighter gray matter. The correct ordering of the gray matter and white matter is achieved by taking the negative of the green channel of the blockface image. Then the background region of the blockface images, mostly consisting of dry ice crystals, was masked out (set to zero intensity) by applying a random forest classifier in ITK-SNAP [84] trained on a small set of user-annotated blockface images. For most blocks (n=48), the classifier trained using a common set of 13 blockface images sampled from across all specimens produced acceptable masking results; however for some blocks (n=25), due to differences in illumination, custom training was performed using a few images from the same specimen. An example of a 3D blockface transformed to MRI-like appearance is in Supplemental Figure S.5b.

##### Registration between Blockface Volume and MRI

The matching between MRI and blockface volumes was initialized manually for each specimen. An ITK-SNAP workspace was created containing the 7T MRI, rotated and translated into the space of the cutting mold model, and the multiple blockface volumes for the specimen. The manual registration tool in ITK-SNAP was used to set the *z*-position of each block and to rotate each block around the *z* axis to achieve a rough alignment. The *z*-positioning of each block is known since the blocks are sectioned according to a 3D printed mold.

The manual registration was used to initialize automatic affine registration, which was performed in the space of each blockface volume. Registration was first performed between the “MRI-like” blockface image and the 7T MRI, and subsequently between the “MRI-like” blockface image and the distortion-corrected 9.4T MRI (which has greater resolution and anatomical detail). Even though the cutting mold reduces the dimensionality of the registration problem by 3 parameters (i.e., position of the block along the mold’s axis is known, and rotation is only unknown around this axis), we perform full 12-parameter affine registration accounting for slight distortion of tissue that may occur. Parameters of this registration are listed in Supplemental Table S.4B, and a sample registration result is shown in Supplemental Figure S.5.

#### 5.5.3. Nissl Slide Intensity Remapping

##### Overview

Direct registration between histology and MRI is challenging because the resolution and intensity characteristics of these two modalities are vastly different. The conventional approach of using information theoretic similarity metrics [54, 67] to register multi-modality images did not perform well in our data, particularly during deformable registration. In our prior work [1], good registration between MRI slices and histology slides stained with the Kluwer-Barrera stain was achieved by downsampling histology to the resolution of the MRI and combining the red, green and blue channels of the downsampled histology images into a monochrome image resembling MRI in appearance. The transformation parameters were derived using least-square fitting between roughly registered histology and corresponding MRI slices. However, this approach does not work as well for the thionin stain because the contrast between gray and white matter is weaker than in the Kluwer-Barrera stain.

In the present study, we take advantage of the fact that at the full resolution level, the characteristics of the thionin/Nissl image at full resolution are very different between gray and white matter regions, as well as between different layers of the cortex. These differences are characterized by the density of neurons vs. glial cells, as well as cell morphology. We use an unsupervised machine learning algorithm to transform full-resolution Nissl histology images into lower-resolution feature images. The full-resolution Nissl image is broken up into non-overlapping regions roughly the size of an MRI voxel, and each region is described by a set of features. These feature images can then be combined linearly with different weights to generate MRI-like appearance.

##### Feature Map Generation from Full-Resolution Nissl Slides

To generate descriptive features from Nissl slides, we make an assumption that patches in the full-resolution slide fall into a finite number of characteristic classes (e.g., white matter, background, layer I of the cortex, granular cell layer of the hippocampus). We used the deep learning based clustering algorithm DeepCluster [22]^1^ to assign Nissl patches selected at random from our archive to a set of *K* = 20 clusters. We sampled a total of 20,480 patches of 256 x 256 pixels at random from across 40 randomly selected Nissl slides in our archive (512 patches per slide) and used 90% of the patches to train DeepCluster. DeepCluster iteratively alternates between a clustering step, in which patches are processed using the AlexNet [46] CNN (initially with random weights) and assigned to *K* classes using the k-means algorithm [49]; and a learning step, in which the AlexNet weights are adjusted by training the network to classify patches into the *K* classes assigned in the clustering step [22]. Once DeepCluster has been trained, applying the AlexNet network to any 256 x 256 pixel patch results in a set of *K* activation values (which can be interpreted as log-likelihoods of the patch belonging to each cluster). We then divide each full-resolution Nissl slide into a grid of non-overlapping 256 x 256 pixel patches, process each patch with the trained AlexNet, and obtain a lower-resolution (by factor of 256) *K*-channel image that implicitly captures the local microscopic patterns in the slide. The clustering and examples of *K*-channel images derived from the Nissl slides are illustrated in Supplemental Figure S.6.

##### Mapping Nissl Sections to MRI-like Appearance

We employ two methods to generate “MRI-like” appearance from the *K*-channel images derived from Nissl slides. The first, “rough”, approach is used before any registration between histology and MRI has taken place. We randomly selected 25 *K*-channel images from across all specimens, and manually labeled examples of gray matter, white matter, and background. We then trained the random forest classifier [20, 26] in ITK-SNAP to discriminate these three tissue classes from the *K*-channel data in the neighborhood of every pixel. This classifier is then applied for every Nissl slide, yielding maps of gray matter, white matter and background probabilities. Lastly, a weighted sum of these probabilities was taken (weighted 0 for background, 100 for white matter, 200 for gray matter) to achieve a MRI-resembling appearance. Examples of this transformation are in Supplemental Figure S.7.

The second, “refined”, approach is employed after initially registering these “rough” MRI-like images to corresponding MRI slices, as described below. After the initial affine registration, most gray matter pixels in the Nissl histology map to gray matter pixels in MRI, and likewise for white matter and background. For each block, we randomly sample 600 pixels from the *K*-channel images derived from the Nissl slides and 600 pixels at the corresponding locations in the MRI slices. We then perform robust linear regression using the rlm function in R [73] with the *K* features as independent variables, MRI intensity as the dependent variable, and the 600 pixels treated as observations. This yields a model of the MRI signal as a linear combination of the *K*-channels, which we then apply to every *K*-channel image in the block. The resulting MRI-like appearance has characteristics strikingly similar to the actual MRI, as illustrated in Supplemental Figure S.7.

#### 5.5.4. Registration between Nissl Histology and MRI

This stage involves performing 2D registration between each Nissl slide and the corresponding slice of the target MRI volume. The target volume, *I_trg_*, is the 9.4T MRI that has been corrected for geometrical distortions (Sec. 5.5.1) and resampled into the space of the blockface volume via affine registration (Sec. 5.5.2). Since each Nissl slide corresponds to a blockface image, the registration between Nissl slides and corresponding slices of *I*_trg_ is a 2D non-linear registration problem. However, performing these 2D registrations independently for every slide would result in highly discontinuous reconstructed histology volumes, since registration errors are unavoidable, and without coordination between neighboring Nissl slides, corresponding locations in adjacent Nissl slides may map to distant locations in *I*_trg_. To overcome this, we employ a multi-step approach, implemented in the open-source tool `stack_greedy`^2^, which builds upon our prior work on histology 3D reconstruction [82, 2, 1]. The approach involves the following stages:

##### Stage 1: Initial Graph-Based Alignment of all Nissl Slides in the Block

Nissl slides are rigidly registered to neighboring Nissl slides up to 1.6 mm away in the *z* dimension (i.e., three adjacent slices on each side). This registration uses the NCC metric applied to the “rough” MRI-like images derived from the Nissl slides. To account for nearly arbitrary orientation of the adjacent Nissl slides, these registrations are initialized using a brute search over the space of rotations and translations. A weighted directed graph is constructed in which the edges represent pairwise registrations, and edge weights encode both the proximity between a pair of neighboring slides in *z* and the quality of the registration between them (derived from the NCC metric). Using Dijkstra’s algorithm, we find shortest paths in this graph, and a root node, for which the sum of shortest paths to all other nodes is smallest. We then compose the rigid transformations along the shortest path from each node to the root node to form a rigid transformation from each node to the root node. This approach, originally introduced in [82], helps ensure that erroneous registrations do not propagate across the reconstructed Nissl volume.

##### Stage 2: Global 2D Affine Matching of Reconstructed Nissl Volume to the Target Volume

This stage finds a global 2D affine transformation in the *x – y* plane that best matches the reconstructed Nissl volume from Stage 1 to *I*_trg_. First, each rough “MRI-like” Nissl slide is independently registered to the corresponding slice of *I*_trg_ using 2D affine registration with the NCC metric. Like in Stage 1, a brute search is performed to initialize these registrations. Among all the individual per-slice affine transformations derived in this way, the one that minimizes the total NCC metric between the reconstructed Nissl volume and *I*_trg_ is selected and applied to each individual Nissl slide. This somewhat exhaustive approach helps ensure consistent initial matching of the histology stack to *I*_trg_ across all blocks.

##### Stage 3: Iterative 2D Affine Matching

In an iterative scheme that passes through the slides in the block 10 times, each Nissl slide is matched to the corresponding slice in *I*_trg_ and to the directly adjacent Nissl slides using affine registration. This step is a refinement upon Stage 2, as it allows slides to transform relative to neighboring slides, while still enforcing consistency across the Nissl stack. The registrations are applied to MRI-like Nissl images and the NCC metric is used. For the first 5 iterations, the “rough” MRI-like Nissl images are used. After the fifth iteration, the “refined” MRI-like images are derived from the Nissl slides (Sec. 5.5.3) and used for the subsequent 5 iterations. After each iteration, an average of the affine transformations from the *i*-th Nissl slide to the corresponding *I*_trg_ slice and to the (*i* – 1)-th and (*i* + 1)-th Nissl slide is computed, and the resulting affine transformation is applied to slide *i* in the following iteration.

##### Stage 4: Iterative 2D Deformable Matching

A similar iterative scheme is carried out for 10 iterations, but now the registrations between Nissl slides, their neighbors, and corresponding *I*_trg_ slices are deformable. The overall effect of this Stage is to correct for local distortions on Nissl slides while maintaining 3D continuity of the slide stack. For each deformable registration, the problem is set up to find a diffeomorphic transformation acting on the *i*-th Nissl slide that minimizes the weighted sum of NCC metrics between Nissl slide *i* and Nissl slide *i* – 1 (with weight 0.25), Nissl slide *i* and Nissl slide *i* + 1 (with weight 0.25) and Nissl slide *i* and the corresponding *I*_trg_ slice (with weight 1.0). The NCC metric is applied to the “refined” MRI-like images. The resulting deformation is applied to slide *i* in subsequent registrations.

Additional details and parameters for these registration steps are in Supplemental Table S.4D,E, and intermediate results for one block are illustrated in Supplemental Figure S.8.

#### 5.5.5. Registration of Tau Slides to Nissl Slides

The IHC slides in this study are lightly counterstained with thionin, making it possible to register them directly to the Nissl slides using the NCC metric. However, during IHC processing, the IHC sections undergo significantly more tearing and displacement of tissue than Nissl slides. As the result, we found that the combination of affine and deformable registration used elsewhere in the paper did not work well for this matching problem.

Instead, we employed an algorithm that takes advantage of the observations that, apart from the tearing and displacement, the tissue undergoes little non-linear deformation, and that tearing tends to follow existing sulcal patterns, ventricular and vascular spaces. To account for tearing, we perform registration in a piecewise manner. A binary tissue/background mask is generated for the Nissl slide, and the METIS graph partitioning algorithm [43] is used to break the mask into a fixed number of contiguous regions (we empirically chose to use 8 regions). When significant tearing is present, different sides of the tear tend to be assigned to different regions, as do different gyri. Affine and deformable registration between the Nissl slide and the IHC slide is performed independently for each of the 8 regions following initial whole-slide rigid alignment with a brute force search initialization. This strategy works surprisingly well for most slides resulting in visually excellent registrations. The piecewise diffeomorphic registrations from the Nissl slide to the IHC slide are used to transform the NFT burden maps into the Nissl slide space. Parameters of this registration strategy are listed in Supplemental Table S.4C, and examples are illustrated in Supplemental Figure S.9.

#### 5.5.6. Evaluation of Registration Accuracy

To evaluate the accuracy of the reconstruction pipeline, Nissl histology and 9.4T MRI were manually annotated in 10 specimens. Four anatomical curves (Supplemental Table S.2) were defined and traced by the same rater (SAL) independently in both modalities. Each curve was drawn on one Nissl slide per specimen. When drawing curves on the MRI, the scan was oriented so that the slicing direction corresponded to that of the histology, and curves were traced on the MRI slice defined in Table S.2, as well as on the four adjacent anterior and four adjacent posterior slices. This was to account for difficulty visually matching MRI and histology in the *z* dimension.

After registration the annotations from the MRI space were resampled into the space of annotated Nissl slides, and distance between the corresponding curves was measured. We computed symmetric root mean squared distance (S-RSMD) between the curves for different stages of the registration pipeline. To account for differences in the extent of the curves, we trimmed away from the ends of the MRI curve all points for which the closed point on the Nissl curve was the endpoint of the Nissl curve; and vice versa. The validation curves are illustrated in Supplemental Figure S.11.

#### 5.5.7. Final 3D Reconstruction

The entire chain of transformations computed in Sec. 5.5.5, 5.5.4 and 5.5.2 is composed and applied to the Nissl slides, IHC slides, and derived NFT burden maps, transforming these 2D images into the space of the distortion-corrected whole-specimen 9.4T MRI. The foreground masks for the Nissl slides are mapped into the distortion-corrected whole-specimen 9.4T MRI space, generating a 3D histology mask volume, which is used to differentiate regions for which NFT burden information is available and regions for which it is missing. Figure 3 shows an example of a whole-specimen reconstruction of Nissl histology, IHC histology, NFT burden map, and histology mask volume, alongside the distortion-corrected 9.4T MRI. Such reconstructions were computed for all specimens in the study and are included in the supplemental digital archive.

#### 5.5.8. Group-Level Analysis of NFT Burden

To allow group-level analysis of three-dimensional NFT burden maps, we warped all the distortion-corrected 9.4T MRI scans into a common anatomical template. This step incorporated *in vivo* 3 Tesla MRI data from the Aging Brain Cohort (ABC) study at the University of Pennsylvania. We first constructed an unbiased population template [42, 9] using the T1-weighted MRI scans of 23 ABC research participants (Supplemental Table S.3). The template captures the average brain anatomy in this population. We then mapped the 3D fluid-attenuated inversion recovery (FLAIR) MRI scans of these participants into the template and averaged them, creating a FLAIR template. The FLAIR template (Supplemental Figure S.12) is more similar to the ex vivo MRI scans than T1 or T2-weighted MRI, with gray matter brighter than white matter, and cerebrospinal fluid and air having the least brightness. The 18 individual 9.4 Tesla MRI scans were linearly and deformably registered to the FLAIR template with the initial rigid transformation performed manually using the ITK-SNAP tool. During this registration, masks in the 9.4T MRI space were used so that edges along which tissue was cut in the 9.4T MRI scan did not contribute to the registration objective function. Once the 9.4T MRI scans were warped into the in vivo MRI space, the unbiased population template algorithm was applied to these images, helping improve groupwise registration quality further. The result of this two-stage template building process was an ex vivo 9.4T MRI template of the MTL spatially matched to the in vivo 3 Tesla MRI template (Supplemental Figure S.12). Parameters the registration steps involved in building the template are listed in Supplemental Table S.4F.

Placing ex vivo information into the in vivo MRI space allows us to visualize NFT burden maps in a familiar space, and to bring in existing annotations and other data available in the in vivo imaging domain. Specifically, in this paper, we bring in manual segmentations of the MTL subregions performed in high-resolution T2-weighted MRI scans of ABC participants using a 3 Tesla MRI protocol derived from [11] into the in vivo template space. We also bring in the anatomical labels from the Human Connectome Project (HCP) Glasser et al. [34] multi-modal atlas into the template space. These anatomical labels allow us to compare NFT burden between anatomical regions of interest (ROIs) without having to manually delineate these ROIs directly in ex vivo MRI space^3^.

## Footnotes

1 Implementation from https://github.com/facebookresearch/deepcluster

2 https://github.com/pyushkevich/greedy

3 We have begun to generate cytoarchitectonically based segmentations of MTL subregion in histology space, but this effort is extremely labor-intensive, and will take years to complete.

## References

[1] D. H. Adler, L. E. M. Wisse, R. Ittyerah, J. B. Pluta, S-L. Ding, L. Xie, J. Wang, S. Kadivar, J. L. Robinson, T. Schuck, J. Q. Trojanowski, M. Grossman, J. A. Detre, M. A. Elliott, J. B. Toledo, W. Liu, S. Pickup, M. I. Miller, S. R. Das, D. A. Wolk, and P. A. Yushkevich. Characterizing the human hippocampus in aging and Alzheimer’s disease using a computational atlas derived from ex vivo MRI and histology. Proc Natl Acad Sci U S A, 115(16):4252–4257, 04 2018. doi: 10.1073/pnas.1801093115.

[2] Daniel H Adler, John Pluta, Salmon Kadivar, Caryne Craige, James C Gee, Brian B Avants, and Paul A Yushkevich. Histology-derived volumetric annotation of the human hippocampal subfields in postmortem MRI. Neuroimage, 84:505–23, Jan 2014. doi: 10.1016/j.neuroimage.2013.08.067.

[3] Sahar Ahmad, Jingfan Fan, Pei Dong, Xiaohuan Cao, Pew-Thian Yap, and Dinggang Shen. Deep learning deformation initialization for rapid groupwise registration of inhomogeneous image populations. Front Neuroinform, 13:34, 2019. doi: 10.3389/fninf.2019.00034.

[4] Maryana Alegro, Yuheng Chen, Dulce Ovando, Helmut Heinser, Rana Eser, Daniela Ushizima, Duygu Tosun, and Lea T Grinberg. Deep learning for Alzheimer’s disease: Mapping large-scale histological tau protein for neuroimaging biomarker validation. 2020. URL https://doi.org/10.1101/698902.

[5] S E Arnold, B T Hyman, J Flory, A R Damasio, and G W Van Hoesen. The topographical and neuroanatomical distribution of neurofibrillary tangles and neuritic plaques in the cerebral cortex of patients with Alzheimer’s disease. Cereb Cortex, 1(1):103–16, 1991. doi: 10.1093/cercor/1.1.103.

[6] S E Arnold, B T Hyman, and G W Van Hoesen. Neuropathologic changes of the temporal pole in Alzheimer’s disease and Pick’s disease. Arch Neurol, 51(2):145–50, Feb 1994. doi: 10.1001/archneur.1994.00540140051014.

[7] Johannes Attems and Kurt A Jellinger. The overlap between vascular disease and alzheimer’s disease–lessons from pathology. BMC Med, 12:206, Nov 2014. doi: 10.1186/s12916-014-0206-2.

[8] B. Avants, C. Epstein, M. Grossman, and J. Gee. Symmetric diffeomorphic image registration with cross-correlation: Evaluating automated labeling of elderly and neurodegenerative brain. Medical Image Analysis, 12:26–41, 2008.

[9] Brian B. Avants and James C. Gee. Shape averaging with diffeomorphic flows for atlas creation. In Proc IEEE Int Symp Biomed Imaging, pages 595–598, 2004.

[10] Alexandre Bejanin, Melissa E Murray, Peter Martin, Hugo Botha, Nirubol Tosakulwong, Christopher G Schwarz, Matthew L Senjem, Gael Chételat, Kejal Kantarci, Clifford R Jack, Bradley F Boeve, David S Knopman, Ronald C Petersen, Caterina Giannini, Joseph E Parisi, Dennis W Dickson, Jennifer L Whitwell, and Keith A Josephs. Antemortem volume loss mirrors tdp-43 staging in older adults with non-frontotemporal lobar degeneration. Brain, 142(11):3621–3635, Nov 2019. doi: 10.1093/brain/awz277.

[11] D Berron, P Vieweg, A Hochkeppler, J B Pluta, S-L Ding, A Maass, A Luther, L Xie, S R Das, D A Wolk, T Wolbers, P A Yushkevich, E Düzel, and L E M Wisse. A protocol for manual segmentation of medial temporal lobe subregions in 7 tesla mri. Neuroimage Clin, 15:466–482, 2017. doi: 10.1016/j.nicl.2017.05.022.

[12] Gabriel Besson, Jessica Simon, Eric Salmon, and Christine Bastin. Familiarity for entities as a sensitive marker of antero-lateral entorhinal atrophy in amnestic mild cognitive impairment. Cortex, 128:61–72, Jul 2020. doi: 10.1016/j.cortex.2020.02.022.

[13] M. Bobinski, J. Wegiel, M. Tarnawski, M. Bobinski, B. Reisberg, M. J. de Leon, D. C. Miller, and H. M. Wisniewski. Relationships between regional neuronal loss and neurofibrillary changes in the hippocampal formation and duration and severity of Alzheimer disease. J Neuropathol Exp Neurol, 56(4):414–420, Apr 1997.

[14] Sethu K Boopathy Jegathambal, Kelvin Mok, David A Rudko, and Amir Shmuel. MRI based brain-specific 3D-printed model aligned to stereotactic space for registering histology to MRI. Annu Int Conf IEEE Eng Med Biol Soc, 2018:802–805, Jul 2018. doi: 10.1109/EMBC.2018.8512346.

[15] H Braak and E Braak. On areas of transition between entorhinal allocortex and temporal isocortex in the human brain. Normal morphology and lamina-specific pathology in Alzheimer’s disease. Acta Neuropathol, 68(4):325–32, 1985. doi: 10.1007/BF00690836.

[16] H. Braak and E. Braak. Neuropathological stageing of Alzheimer-related changes. Acta Neuropathol, 82(4):239–259, 1991.

[17] H. Braak and E. Braak. Staging of Alzheimer’s disease-related neurofibrillary changes. Neurobiol Aging, 16(3):271–8; discussion 278–84, 1995.

[18] H Braak and E Braak. Frequency of stages of alzheimer-related lesions in different age categories. Neurobiol Aging, 18(4):351–7, 1997.

[19] Heiko Braak, Dietmar R Thal, Estifanos Ghebremedhin, and Kelly Del Tredici. Stages of the pathologic process in alzheimer disease: age categories from 1 to 100 years. J Neuropathol Exp Neurol, 70(11):960–9, Nov 2011. doi: 10.1097/NEN.0b013e318232a379.

[20] Leo Breiman. Random forests. Machine learning, 45(1):5–32, 2001.

[21] Johannes Brettschneider, Kelly Del Tredici, Virginia M-Y Lee, and John Q Trojanowski. Spreading of pathology in neurodegenerative diseases: a focus on human studies. Nat Rev Neurosci, 16(2):109–20, Feb 2015. doi: 10.1038/nrn3887.

[22] Mathilde Caron, Piotr Bojanowski, Armand Joulin, and Matthijs Douze. Deep clustering for unsupervised learning of visual features. In Proceedings of the European Conference on Computer Vision (ECCV), pages 132–149, 2018.

[23] Paolo Cignoni, Massimiliano Corsini, and Guido Ranzuglia. Meshlab: an open-source 3D mesh processing system. ERCIM News, 2008(73), 2008. URL http://dblp.uni-trier.de/db/journals/ercim/ercim2008.html#CignoniCR08.

[24] Martí Colom-Cadena, Ellen Gelpi, Sara Charif, Olivia Belbin, Rafael Blesa, Maria J Martí, Jordi Clarimón, and Alberto Lleó. Confluence of α-synuclein, tau, and β-amyloid pathologies in dementia with Lewy bodies. J Neuropathol Exp Neurol, 72(12):1203–12, Dec 2013. doi: 10.1097/NEN.0000000000000018.

[25] John F Crary, John Q Trojanowski, Julie A Schneider, Jose F Abisambra, Erin L Abner, Irina Alafuzoff, Steven E Arnold, Johannes Attems, Thomas G Beach, Eileen H Bigio, Nigel J Cairns, Dennis W Dickson, Marla Gearing, Lea T Grinberg, Patrick R Hof, Bradley T Hyman, Kurt Jellinger, Gregory A Jicha, Gabor G Kovacs, David S Knopman, Julia Kofler, Walter A Kukull, Ian R Mackenzie, Eliezer Masliah, Ann McKee, Thomas J Montine, Melissa E Murray, Janna H Neltner, Ismael Santa-Maria, William W Seeley, Alberto Serrano-Pozo, Michael L Shelanski, Thor Stein, Masaki Takao, Dietmar R Thal, Jonathan B Toledo, Juan C Troncoso, Jean Paul Vonsattel, Charles L White, 3rd, Thomas Wisniewski, Randall L Woltjer, Masahito Yamada, and Peter T Nelson. Primary age-related tauopathy (part): a common pathology associated with human aging. Acta Neuropathol, 128(6):755–66, Dec 2014. doi: 10.1007/s00401-014-1349-0.

[26] Antonio Criminisi, Jamie Shotton, and Ender Konukoglu. Decision forests: A unified framework for classification, regression, density estimation, manifold learning and semi-supervised learning. Foundations and Trends in Computer Graphics and Vision, 7(2–3):81–227, 2012.

[27] Sandhitsu R Das, Lauren Mancuso, Ingrid R Olson, Steven E Arnold, and David A Wolk. Short-term memory depends on dissociable medial temporal lobe regions in amnestic mild cognitive impairment. Cereb Cortex, 26(5):2006–17, May 2016. doi: 10.1093/cercor/bhv022.

[28] Robin de Flores, Laura E M Wisse, Sandhitsu R Das, Long Xie, Corey T McMillan, John Q Trojanowski, John L Robinson, Murray Grossman, Edward Lee, David J Irwin, Paul A Yushkevich, and David A Wolk. Contribution of mixed pathology to medial temporal lobe atrophy in Alzheimer’s disease. Alzheimers Dement, 16(6):843–852, 06 2020. doi: 10.1002/alz.12079.

[29] Dennis W Dickson. Neuropathology of non-alzheimer degenerative disorders. Int J Clin Exp Pathol, 3(1):1–23, 2009.

[30] Song-Lin Ding, Gary W Van Hoesen, Martin D Cassell, and Amy Poremba. Parcellation of human temporal polar cortex: a combined analysis of multiple cytoarchitectonic, chemoarchitectonic, and pathological markers. J Comp Neurol, 514(6):595–623, Jun 2009. doi: 10.1002/cne.22053.

[31] Thibaut Durand, Taylor Mordan, Nicolas Thome, and Matthieu Cord. Wildcat: Weakly supervised learning of deep convnets for image classification, pointwise localization and segmentation. In Proceedings of the IEEE conference on computer vision and pattern recognition, pages 642–651, 2017.

[32] Garrett S Gibbons, Rachel A Banks, Bumjin Kim, Lakshmi Changolkar, Dawn M Riddle, Susan N Leight, David J Irwin, John Q Trojanowski, and Virginia M Y Lee. Detection of Alzheimer disease (AD)-specific tau pathology in AD and nonAD tauopathies by immunohistochemistry with novel conformation-selective tau antibodies. J Neuropathol Exp Neurol, 77(3):216–228, 03 2018. doi: 10.1093/jnen/nly010.

[33] Garrett S Gibbons, Soo-Jung Kim, John L Robinson, Lakshmi Changolkar, David J Irwin, Leslie M Shaw, Virginia M-Y Lee, and John Q Trojanowski. Detection of Alzheimer’s disease (AD) specific tau pathology with conformation-selective anti-tau monoclonal antibody in co-morbid frontotemporal lobar degeneration-tau (FTLD-tau). Acta Neuropathol Commun, 7(1):34, 03 2019. doi: 10.1186/s40478-019-0687-5.

[34] Matthew F Glasser, Timothy S Coalson, Emma C Robinson, Carl D Hacker, John Harwell, Essa Yacoub, Kamil Ugurbil, Jesper Andersson, Christian F Beckmann, Mark Jenkinson, Stephen M Smith, and David C Van Essen. A multi-modal parcellation of human cerebral cortex. Nature, 536(7615):171–178, 08 2016. doi: 10.1038/nature18933.

[35] K. He, X. Zhang, S. Ren, and J. Sun. Deep residual learning for image recognition. In Proceedings of the IEEE conference on computer vision and pattern recognition, pages 770–778, 2016.

[36] B. Hyman, C. Phelps, T. Beach, E. Bigio, N. Cairns, M. Carrillo, D. Dickson, C. Duyckaerts, M. Frosch, E. Masliah, S. Mirra, P. Nelson, J. Schneider, D. Thal, B. Thies, J. Trojanowski, H. Vinters, and T. Montine. National Institute on Aging-Alzheimer’s Association guidelines for the neuropathologic assessment of Alzheimer’s disease. Alzheimers Dement, 8(1):1–13, Jan 2012. doi: 10.1016/j.jalz.2011.10.007.

[37] Juan Eugenio Iglesias, Marc Modat, Loïc Peter, Allison Stevens, Roberto Annunziata, Tom Vercauteren, Ed Lein, Bruce Fischl, Sebastien Ourselin, and Alzheimer’s Disease Neuroimaging Initiative. Joint registration and synthesis using a probabilistic model for alignment of MRI and histological sections. Med Image Anal, 50:127–144, 12 2018. doi: 10.1016/j.media.2018.09.002.

[38] Clifford R Jack, Jr, David A Bennett, Kaj Blennow, Maria C Carrillo, Billy Dunn, Samantha Budd Haeberlein, David M Holtzman, William Jagust, Frank Jessen, Jason Karlawish, Enchi Liu, Jose Luis Molinuevo, Thomas Montine, Creighton Phelps, Katherine P Rankin, Christopher C Rowe, Philip Scheltens, Eric Siemers, Heather M Snyder, Reisa Sperling, and Contributors. Nia-aa research framework: Toward a biological definition of alzheimer’s disease. Alzheimers Dement, 14(4):535–562, 04 2018. doi: 10.1016/j.jalz.2018.02.018.

[39] William Jagust. Imaging the evolution and pathophysiology of alzheimer disease. Nat Rev Neurosci, 19(11):687–700, 11 2018. doi: 10.1038/s41583-018-0067-3.

[40] Bryan D James, Robert S Wilson, Patricia A Boyle, John Q Trojanowski, David A Bennett, and Julie A Schneider. TDP-43 stage, mixed pathologies, and clinical Alzheimer’s-type dementia. Brain, 139(11):2983–2993, 11 2016. doi: 10.1093/brain/aww224.

[41] Keith A Josephs, Melissa E Murray, Jennifer L Whitwell, Nirubol Tosakulwong, Stephen D Weigand, Leonard Petrucelli, Amanda M Liesinger, Ronald C Petersen, Joseph E Parisi, and Dennis W Dickson. Updated tdp-43 in alzheimer’s disease staging scheme. Acta Neuropathol, 131(4):571–85, Apr 2016. doi: 10.1007/s00401-016-1537-1.

[42] S. Joshi, Brad Davis, Matthieu Jomier, and Guido Gerig. Unbiased diffeomorphic atlas construction for computational anatomy. Neuroimage, 23 Suppl 1:S151–S160, 2004. doi: 68.

[43] G. Karypis and V. Kumar. METIS – a software package for partitioning unstructured graphs, partitioning meshes, and computing fill-reducing orderings of sparse matrices – version 4.0. University of Minnesota, 1998.

[44] Sasa L Kivisaari, Lorraine K Tyler, Andreas U Monsch, and Kirsten I Taylor. Medial perirhinal cortex disambiguates confusable objects. Brain, 135(Pt 12):3757–69, Dec 2012. doi: 10.1093/brain/aws277.

[45] Gabor G Kovacs, Sharon X Xie, John L Robinson, Edward B Lee, Douglas H Smith, Theresa Schuck, Virginia M-Y Lee, and John Q Trojanowski. Sequential stages and distribution patterns of aging-related tau astrogliopathy (artag) in the human brain. Acta Neuropathol Commun, 6(1):50, 06 2018. doi: 10.1186/s40478-018-0552-y.

[46] Alex Krizhevsky, Ilya Sutskever, and Geoffrey E Hinton. Imagenet classification with deep convolutional neural networks. In Advances in neural information processing systems, pages 1097–1105, 2012.

[47] L J Kromer Vogt, B T Hyman, G W Van Hoesen, and A R Damasio. Pathological alterations in the amygdala in Alzheimer’s disease. Neuroscience, 37(2):377–85, 1990. doi: 10.1016/0306-4522(90)90408-v.

[48] John Lafferty, Andrew McCallum, and Fernando CN Pereira. Conditional random fields: Probabilistic models for segmenting and labeling sequence data. 2001.

[49] Stuart Lloyd. Least squares quantization in PCM. IEEE transactions on information theory, 28(2):129–137, 1982.

[50] Val J Lowe, Heather J Wiste, Matthew L Senjem, Stephen D Weigand, Terry M Therneau, Bradley F Boeve, Keith A Josephs, Ping Fang, Mukesh K Pandey, Melissa E Murray, Kejal Kantarci, David T Jones, Prashanthi Vemuri, Jonathan Graff-Radford, Christopher G Schwarz, Mary M Machulda, Michelle M Mielke, Rosebud O Roberts, David S Knopman, Ronald C Petersen, and Clifford R Jack, Jr. Widespread brain tau and its association with ageing, Braak stage and Alzheimer’s dementia. Brain, 141(1):271–287, 01 2018. doi: 10.1093/brain/awx320.

[51] Nicholas J Luciano, Pascal Sati, Govind Nair, Joseph R Guy, Seung-Kwon Ha, Martina Absinta, Wen-Yang Chiang, Emily C Leibovitch, Steven Jacobson, Afonso C Silva, and Daniel S Reich. Utilizing 3D printing technology to merge MRI with histology: A protocol for brain sectioning. J Vis Exp, (118), 12 2016. doi: 10.3791/54780.

[52] Anne Maass, David Berron, Theresa M Harrison, Jenna N Adams, Renaud La Joie, Suzanne Baker, Taylor Mellinger, Rachel K Bell, Kaitlin Swinnerton, Ben Inglis, Gil D Rabinovici, Emrah Düzel, and William J Jagust. Alzheimer’s pathology targets distinct memory networks in the ageing brain. Brain, 142(8):2492–2509, 08 2019. doi: 10.1093/brain/awz154.

[53] Ian R A Mackenzie, Matt Baker, Stuart Pickering-Brown, Ging-Yuek R Hsiung, Caroline Lindholm, Emily Dwosh, Jennifer Gass, Ashley Cannon, Rosa Rademakers, Mike Hutton, and Howard H Feldman. The neuropathology of frontotemporal lobar degeneration caused by mutations in the progranulin gene. Brain, 129(Pt 11):3081–90, Nov 2006. doi: 10.1093/brain/awl271.

[54] F. Maes, A. Collignon, D. Vandermeulen, G. Marchal, and P. Suetens. Multi-modality image registration by maximization of mutual information. IEEE Transactions on Medical Imaging, 16(2):187–198, Apr 1997.

[55] Radoslav Matej, Adam Tesar, and Robert Rusina. Alzheimer’s disease and other neurodegenerative dementias in comorbidity: A clinical and neuropathological overview. Clin Biochem, 73:26–31, Nov 2019. doi: 10.1016/j.clinbiochem.2019.08.005.

[56] Jonas Pichat, Eugenio Iglesias, Sotiris Nousias, Tarek Yousry, Sebastien Ourselin, and Marc Modat. Part-to-whole registration of histology and MRI using shape elements. In Proceedings of the IEEE International Conference on Computer Vision (ICCV) Workshops, Oct 2017.

[57] Joseph L Price. Comparative aspects of amygdala connectivity. Ann N Y Acad Sci, 985:50–8, Apr 2003. doi: 10.1111/j.1749-6632.2003.tb07070.x.

[58] Charan Ranganath and Maureen Ritchey. Two cortical systems for memory-guided behaviour. Nat Rev Neurosci, 13(10):713–26, Oct 2012. doi: 10.1038/nrn3338.

[59] S. Ravikumar, L. E. M. Wisse, R. Ittyerah, S. Lim, M. Lavery, L. Xie, J. L. Robinson, T. Schuck, M. Grossman, E. B. Lee, M. D. Tisdall, K. Prabhakaran, J. A. Detre, S. R. Das, G. Mizsei, E. Artacho-Pérula, M. M. I. de Onzono Martin, M. del Mar Arroyo Jiménez, M. Muñoz, F. J. M. Romero, M. del Pilar Marcos Rabal, D. J. Irwin, J. Q. Trojanowski, D. A. Wolk, R. Insausti, and P. A. Yushkevich. Building an ex vivo atlas of the earliest brain regions affected by Alzheimer’s disease pathology. In 2020 IEEE 17th International Symposium on Biomedical Imaging (ISBI), pages 113–117, 2020.

[60] John L Robinson, Edward B Lee, Sharon X Xie, Lior Rennert, EunRan Suh, Colin Bredenberg, Carrie Caswell, Vivianna M Van Deerlin, Ning Yan, Ahmed Yousef, Howard I Hurtig, Andrew Siderowf, Murray Grossman, Corey T McMillan, Bruce Miller, John E Duda, David J Irwin, David Wolk, Lauren Elman, Leo McCluskey, Alice Chen-Plotkin, Daniel Weintraub, Steven E Arnold, Johannes Brettschneider, Virginia M-Y Lee, and John Q Trojanowski. Neurodegenerative disease concomitant proteinopathies are prevalent, age-related and APOE4-associated. Brain, 141(7):2181–2193, 07 2018. doi: 10.1093/brain/awy146.

[61] John L Robinson, Ning Yan, Carrie Caswell, Sharon X Xie, EunRan Suh, Vivianna M Van Deerlin, Garrett Gibbons, David J Irwin, Murray Grossman, Edward B Lee, Virginia M-Y Lee, Bruce Miller, and John Q Trojanowski. Primary tau pathology, not copathology, correlates with clinical symptoms in PSP and CBD. J Neuropathol Exp Neurol, 79(3):296–304, 03 2020. doi: 10.1093/jnen/nlz141.

[62] O. Ronneberger, P. Fischer, and T. Brox. U-net: Convolutional networks for biomedical image segmentation. In International Conference on Medical image computing and computer-assisted intervention, pages 234–241. Springer, 2015.

[63] Julie A Schneider, Patricia A Boyle, Zoe Arvanitakis, Julia L Bienias, and David A Bennett. Subcortical infarcts, Alzheimer’s disease pathology, and memory function in older persons. Ann Neurol, 62(1):59–66, Jul 2007. doi: 10.1002/ana.21142.

[64] Adam J Schwarz, Peng Yu, Bradley B Miller, Sergey Shcherbinin, James Dickson, Michael Navitsky, Abhinay D Joshi, Michael D Devous, Sr, and Mark S Mintun. Regional profiles of the candidate tau PET ligand 18F-AV-1451 recapitulate key features of Braak histopathological stages. Brain, 139(Pt 5):1539–50, 05 2016. doi: 10.1093/brain/aww023.

[65] Maxim Signaevsky, Marcel Prastawa, Kurt Farrell, Nabil Tabish, Elena Baldwin, Natalia Han, Megan A Iida, John Koll, Clare Bryce, Dushyant Purohit, Vahram Haroutunian, Ann C McKee, Thor D Stein, Charles L White, 3rd, Jamie Walker, Timothy E Richardson, Russell Hanson, Michael J Donovan, Carlos Cordon-Cardo, Jack Zeineh, Gerardo Fernandez, and John F Crary. Artificial intelligence in neuropathology: deep learning-based assessment of tauopathy. Lab Invest, 99(7):1019–1029, 07 2019. doi: 10.1038/s41374-019-0202-4.

[66] S.A. Small, S.A. Schobel, R.B. Buxton, M.P. Witter, and C.A. Barnes. A pathophysiological framework of hippocampal dysfunction in ageing and disease. Nature Reviews Neuroscience, 12(10):585–601, 2011.

[67] Colin Studholme, Derek LG Hill, and David J Hawkes. An overlap invariant entropy measure of 3d medical image alignment. Pattern recognition, 32(1):71–86, 1999.

[68] Susan M Sunkin, Lydia Ng, Chris Lau, Tim Dolbeare, Terri L Gilbert, Carol L Thompson, Michael Hawrylycz, and Chinh Dang. Allen brain atlas: an integrated spatio-temporal portal for exploring the central nervous system. Nucleic Acids Res, 41(Database issue):D996–D1008, Jan 2013. doi: 10.1093/nar/gks1042.

[69] Ziqi Tang, Kangway V Chuang, Charles DeCarli, Lee-Way Jin, Laurel Beckett, Michael J Keiser, and Brittany N Dugger. Interpretable classification of Alzheimer’s disease pathologies with a convolutional neural network pipeline. Nat Commun, 10(1):2173, 05 2019. doi: 10.1038/s41467-019-10212-1.

[70] Jon B Toledo, Steven E Arnold, Kevin Raible, Johannes Brettschneider, Sharon X Xie, Murray Grossman, Sarah E Monsell, Walter A Kukull, and John Q Trojanowski. Contribution of cerebrovascular disease in autopsy confirmed neurodegenerative disease cases in the National Alzheimer’s Coordinating Centre. Brain, 136(Pt 9):2697–706, Sep 2013. doi: 10.1093/brain/awt188.

[71] Nicholas J Tustison, Brian B Avants, Philip A Cook, Yuanjie Zheng, Alexander Egan, Paul A Yushkevich, and James C Gee. N4ITK: improved N3 bias correction. IEEE Trans Med Imaging, 29(6):1310–1320, Jun 2010. doi: 10.1109/TMI.2010.2046908. URL http://dx.doi.org/10.1109/TMI.2010.2046908.

[72] Daniel Tward, Timothy Brown, Yusuke Kageyama, Jaymin Patel, Zhipeng Hou, Susumu Mori, Marilyn Albert, Juan Troncoso, and Michael Miller. Diffeomorphic registration with intensity transformation and missing data: Application to 3D digital pathology of Alzheimer’s disease. Front Neurosci, 14:52, 2020. doi: 10.3389/fnins.2020.00052.

[73] William N Venables and Brian D Ripley. Modern applied statistics with S-PLUS. Springer Science & Business Media, 2013.

[74] Tom Vercauteren, Xavier Pennec, Aymeric Perchant, and Nicholas Ayache. Diffeomorphic demons: efficient non-parametric image registration. Neuroimage, 45(1 Suppl):S61–72, Mar 2009. doi: 10.1016/j.neuroimage.2008.10.040.

[75] Lauren Walker, Kirsty E McAleese, Alan J Thomas, Mary Johnson, Carmen Martin-Ruiz, Craig Parker, Sean J Colloby, Kurt Jellinger, and Johannes Attems. Neuropathologically mixed alzheimer’s and lewy body disease: burden of pathological protein aggregates differs between clinical phenotypes. Acta Neuropathol, 129(5):729–48, May 2015. doi: 10.1007/s00401-015-1406-3.

[76] David A Wolk, Lauren Mancuso, Daria Kliot, Steven E Arnold, and Bradford C Dickerson. Familiarity-based memory as an early cognitive marker of preclinical and prodromal AD. Neuropsychologia, 51(6):1094–102, May 2013. doi: 10.1016/j.neuropsychologia.2013.02.014.

[77] David A Wolk, Sandhitsu R Das, Susanne G Mueller, Michael W Weiner, Paul A Yushkevich, and Alzheimer’s Disease Neuroimaging Initiative. Medial temporal lobe subregional morphometry using high resolution mri in alzheimer’s disease. Neurobiol Aging, 49:204–213, 01 2017. doi: 10.1016/j.neurobiolaging.2016.09.011.

[78] Alexander Wurts, Derek H Oakley, Bradley T Hyman, and Siddharth Samsi. Segmentation of tau stained alzheimers brain tissue using convolutional neural networks. Annu Int Conf IEEE Eng Med Biol Soc, 2020:1420–1423, 07 2020. doi: 10.1109/EMBC44109.2020.9175832.

[79] Long Xie, Laura E M Wisse, Das Sandhitsu, Nicolas Vergnet, Mengjin Dong, Ranjit Ittyerah, Robin deFlores, Paul A Yushkevich, and David A Wolk. Longitudinal atrophy in early Braak regions in preclinical Alzheimer’s disease. Human Brain Mapping, 2020. in press.

[80] Xin Yi, Ekta Walia, and Paul Babyn. Generative adversarial network in medical imaging: A review. Med Image Anal, 58:101552, 12 2019. doi: 10.1016/j.media.2019.101552.

[81] P. A. Yushkevich, H. Wang, J. Pluta, and B. B. Avants. From label fusion to correspondence fusion: a new approach to unbiased groupwise registration. In IEEE International Conference on Computer Vision and Pattern Recognition, 2012.

[82] Paul A. Yushkevich, Brian B. Avants, Lydia Ng, Michael Hawrylycz, Pablo D. Burstein, Hui Zhang, and James C. Gee. 3D mouse brain reconstruction from histology using a coarse-to-fine approach. In Workshop on Biomedical Image Registration (WBIR), 2006.

[83] Paul A Yushkevich, John B Pluta, Hongzhi Wang, Long Xie, Song-Lin Ding, Eske C Gertje, Lauren Mancuso, Daria Kliot, Sandhitsu R Das, and David A Wolk. Automated volumetry and regional thickness analysis of hippocampal subfields and medial temporal cortical structures in mild cognitive impairment. Hum Brain Mapp, 36(1):258–87, Jan 2015. doi: 10.1002/hbm.22627.

[84] Paul A Yushkevich, Artem Pashchinskiy, Ipek Oguz, Suyash Mohan, J Eric Schmitt, Joel M Stein, Dženan Zukić, Jared Vicory, Matthew McCormick, Natalie Yushkevich, Nadav Schwartz, Yang Gao, and Guido Gerig. User-guided segmentation of multi-modality medical imaging datasets with itk-snap. Neuroinformatics, 17(1):83–102, 01 2019. doi: 10.1007/s12021-018-9385-x.

