## Supplemental Material for "3D Mapping of Neurofibrillary Tangle Burden in the Human Medial Temporal Lobe"

### Appendix A. Supplemental Methods

#### *Appendix A.1. 3D printing of custom cutting molds*

To generate a custom printing mold from a 7T MRI scan of the MTL, the following steps are taken:

1. The tissue in the 7T MRI scan is segmented from the background using the semi-automatic algorithm in ITK-SNAP [84], as illustrated in Figure S.1a.
2. A template mold image is loaded in ITK-SNAP (purple and beige in Figure S.1b) and the 7T MRI (grayscale in Figure S.1b) is overlaid. The 7T MRI is rotated relative to the mold template to ensure that the hippocampal main axis is aligned with the long axis of the mold and the tissue is enclosed in the mold template.
3. The rotated MTL segmentation of the tissue is extruded upward through the mold template, i.e., every voxel in the mold template that is located above a foreground voxel in the MTL segmentation is set to background intensity.
4. An additional cropping mask is drawn using the ITK-SNAP polygon tool and used to trim the mold image. The crop mask is used to reduce print time and filament use by only printing a narrow shell around the MTL.
5. A 3D model is generated from the mold image and simplified using the quadric edge collapse decimation algorithm in MeshLab software [23]. The model is sent for 3D printing on the Ultimaker 3+ Extended printer (Ultimaker B.V., Utrecht, The Netherlands).
6. A visualization of the 3D mold model and the tissue in intended orientation in the model is generated and printed in color. This visualization is used to guide the placement of actual tissue in the mold (Figure S.1c).

### Appendix B. Supplemental Results

#### *Appendix B.1. MRI and histology registration accuracy at different stages of the registration pipeline*

Supplemental Figure S.10 reports the symmetric root mean squared distance (RMSD) for curves C1-C4 defined in Figure 4 at three stages of the MRI-Nissl registration pipeline (Section 5.5.4): global affine alignment, iterative affine alignment, and iterative deformable alignment. Supplemental figure S.11 shows the alignment for the four median-performance examples in Figure 4) at these three registration stages. The average mismatch is lower for the iterative phases than for the global phase. For curves C2 and C3, the mismatch is substantially lower in the deformable stage than in the iterative affine stage, while for C1 and C4, there is a very small gain from the deformable stage.

### Appendix C. Supplemental Figures and Tables

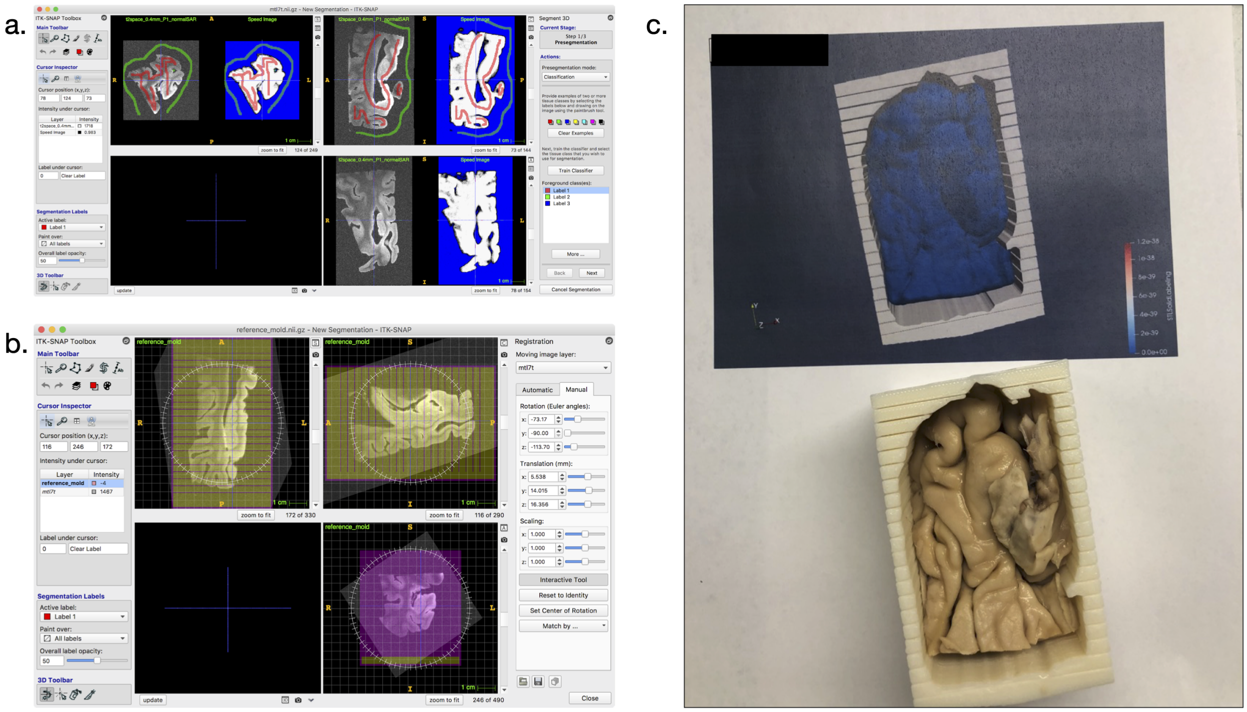

Figure S.1: Steps in the 3D printing pipeline. (a). Segmentation of the MTL from 7T MRI. (b). Orientation of the 7T image (grayscale) relative to the template of the cutting mold. (c). A 3D rendering of the tissue model (blue) in the mold (white) is printed on paper and used by technologists to guide placement of tissue in the mold.

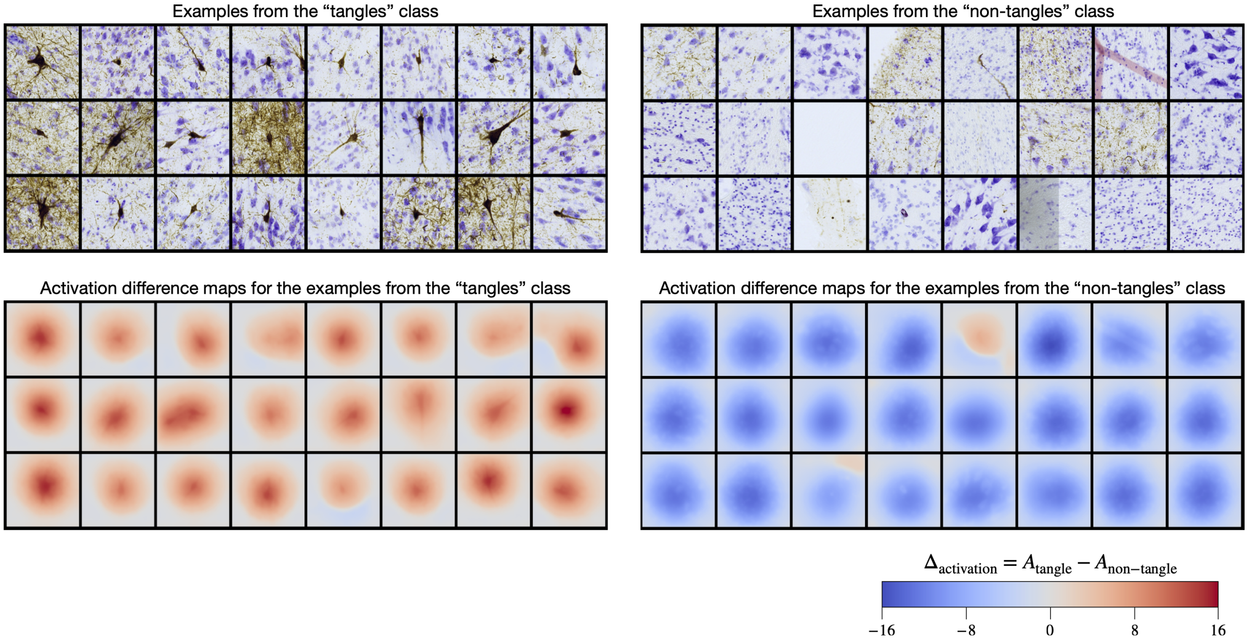

Figure S.2: Examples of patches used to train and evaluate the weakly supervised deep learning algorithm WildCat [31] for NFT burden quantification. Examples of 512x512 pixel patches containing tangle-like and non-tangle objects in the anti-tau immunohistochemistry images are shown at the top. At the bottom, for each patch, the difference between the 112x112 pixel activation map for the “tangle” class and the activation map for the “non-tangle” class is shown. The activation map is a heat map extracted from the last layer of the WildCat network that is pooled spatially and used to assign each patch a class label. Positive values in the activation difference images correspond to the tangle class and negative values to the non-tangle class.

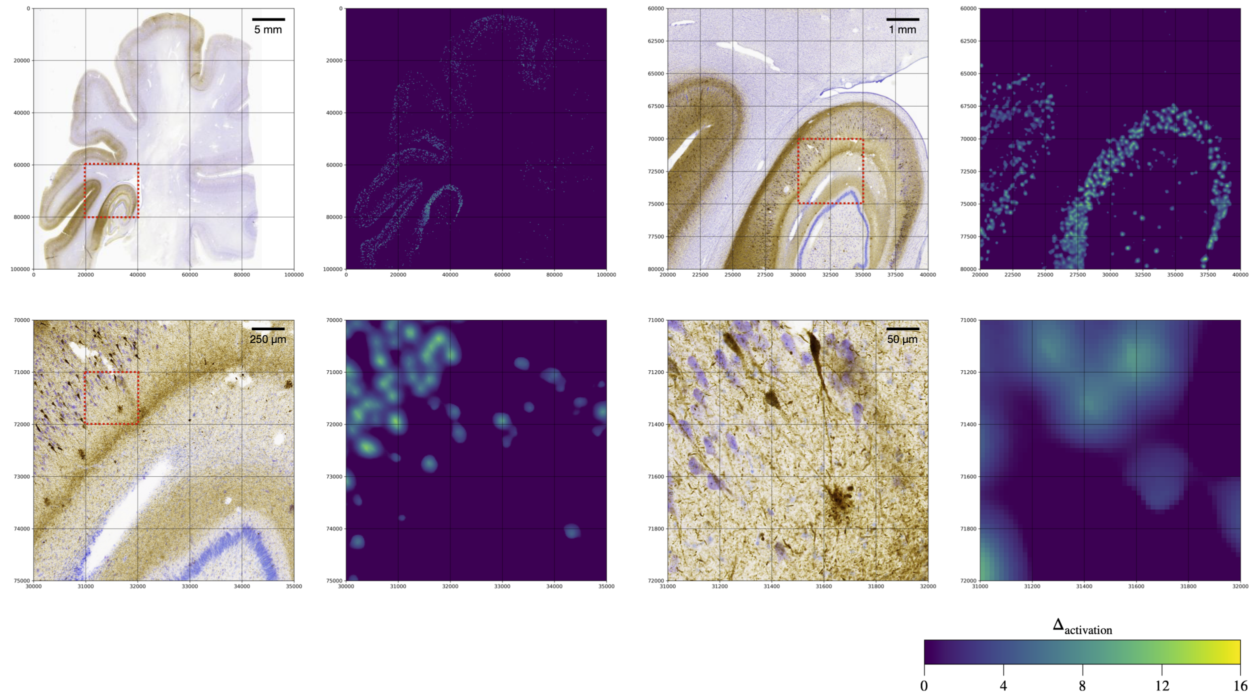

Figure S.3: Example of NFT burden quantification on a whole-slide anti-tau immunohistochemistry image. For four nested regions of interest with increasing level of magnification (indicated a the red box), the immunohistochemistry image and the activation difference map from the WildCat [31] model are shown. The activation difference map is thresholded at zero. NFT burden maps plotted in Figures 5 and 6 downsampled from this thresholded activation difference map.

Table S.1: Extended description of the brain donor cohort, primary and secondary postmortem diagnoses, and global neuropathological staging using the Hyman et al. [36] protocol. Staging and diagnoses were derived from the opposite hemisphere from the one scanned with MRI. Abbreviations: HNL: Human Neuroanatomy Laboratory (University of Castilla-La Mancha at Albacete, Spain); CNDR: Center for Neurodegenerative Disease Research (University of Pennsylvania, USA); PART: probable primary age-related tauopathy; ADNC: Alzheimer's disease neuropathological change; CBD: corticobasal degeneration; LBD: Lewy body disease; AGD: argyrophilic grain disease; FTLD: frontotemporal lobar degeneration of TDP-43 type; PPA: primary progressive aphasia; bvFTD: behavioral variant of frontotemporal dementia; PD: Parkinson's disease.

| | | | | Pathologic Diagnosis | | Global Pathology Score (NIA/AA Criteria) | | | | | |
|---|---|---|---|---|---|---|---|---|---|---|---|
| Participant | Site | Age | Sex | Primary | Secondary | Amyloid (A) | Braak (B) | CERAD (C) | $\alpha$-synuclein | TDP-43 | Clinical Diagnosis |
| 1 | HNL | 45 | Male | PART | | A0 | B1 | C0 | 0 | 0 | N/A |
| 2 | HNL | 76 | Female | Low ADNC | | A1 | B1 | C0 | 0 | 0 | N/A |
| 3 | HNL | 78 | Female | Low ADNC | CVD | A1 | B1 | C1 | 0 | 0 | N/A |
| 4 | HNL | 93 | Male | Intermediate ADNC | | A3 | B2 | C3 | 0 | 0 | N/A |
| 5 | HNL | 71 | Male | Low ADNC | | A1 | B1 | C0 | 0 | 0 | N/A |
| 6 | HNL | 74 | Male | PART | | A0 | B2 | C0 | 0 | 0 | N/A |
| 7 | HNL | 61 | Male | PART | CBD | A0 | B1 | C0 | 0 | 0 | N/A |
| 8 | HNL | 70 | Male | Low ADNC | LBD | A1 | B1 | C0 | 2 | 0 | N/A |
| 9 | HNL | 90 | Male | Low ADNC | | A1 | B1 | C0 | 0 | 0 | N/A |
| 10 | HNL | 76 | Female | Unremarkable | | A0 | B0 | C0 | 0 | 0 | N/A |
| 11 | HNL | 66 | Female | PART | LBD | A0 | B1 | C0 | 1 | 0 | N/A |
| 12 | HNL | 93 | Female | Low ADNC | | A1 | B1 | C1 | 0 | 0 | N/A |
| 13 | CNDR | 76 | Female | CBD | | A1 | B0 | C1 | 0 | 0 | FTLD-PPA |
| 14 | CNDR | 70 | Male | LBD | | A0 | B2 | C0 | 1 | 0 | PD |
| 15 | CNDR | 80 | Male | AGD | PSP | A1 | unratable | C0 | 0 | 0 | FTLD-bvFTD |
| 16 | CNDR | 77 | Male | CBD | FTLD-TDP | A1 | B0 | C1 | 0 | 3 | Corticobasal Syndrome |
| 17 | CNDR | 82 | Female | FTLD-TDP | | A0 | B2 | C0 | 0 | 3 | FTLD-PPA (PNFA) |
| 18 | CNDR | 76 | Male | CVD | Low ADNC | A1 | B1 | C0 | 0 | 0 | Vascular Dementia |

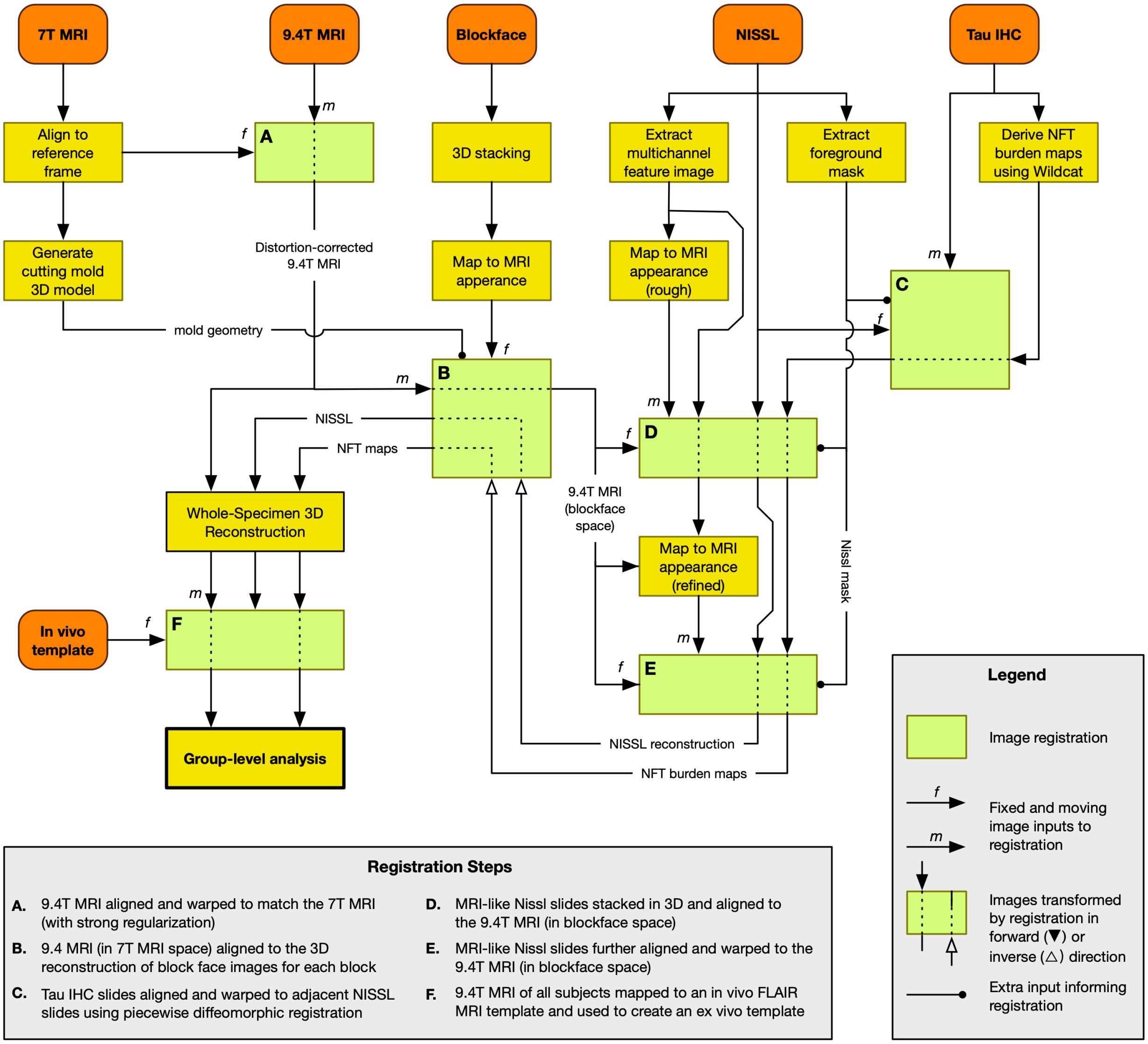

Figure S.4: Flowchart of the complete image analysis pipeline. Five modalities (7T MRI, 9.4T MRI, blockface images, Nissl histology, and anti-tau IHC; examples of each modality shown in Figure 1) are input into a complex pipeline with multiple image analysis and image registration steps. For representing image registrations, we use a special notation (see Legend), where a green box represents a registration; the fixed (or reference) image and the moving image are indicated by arrows with letters 'f' and 'm'; and the action of applying a transformation computed by the registration (affine transformation, deformable warp, or an inverse thereof) is indicated by a dashed line inside the green box.

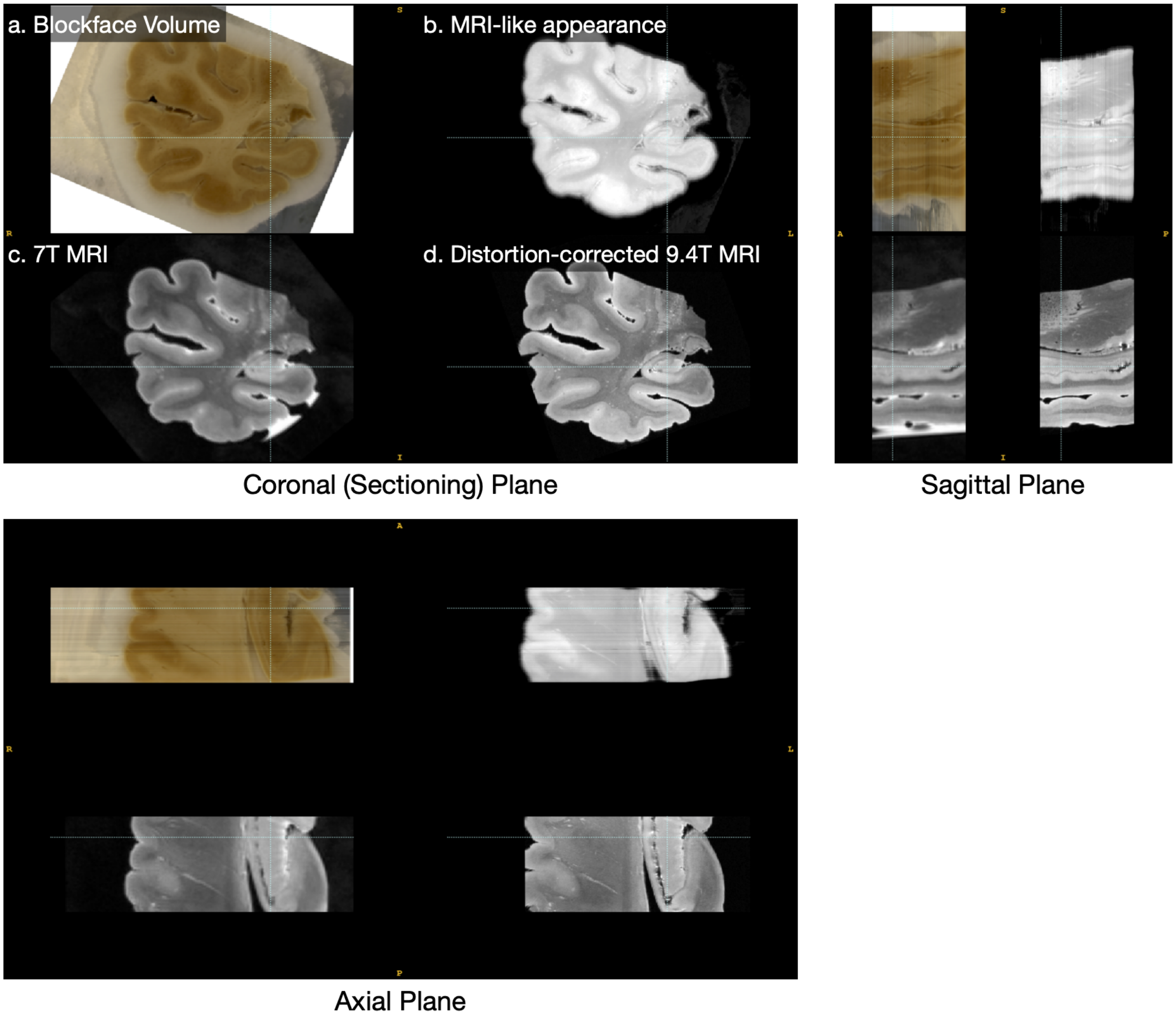

Figure S.5: Blockface image reconstruction and matching to MRI. The blockface images are stacked in 3D with uniform 50µm spacing to form a blockface volume. Using a random forest classifier to separate tissue from background (dry ice) and taking the negative of the green channel of the blockface image yields a grayscale image with roughly MRI-like appearance, which can then be registered with the MRI image using conventional image similarity metrics. The result shown here is from the last stage of blockface-MRI registration, i.e., affine registration between the blockface volume and the distortion-corrected 9.4T MRI volume.

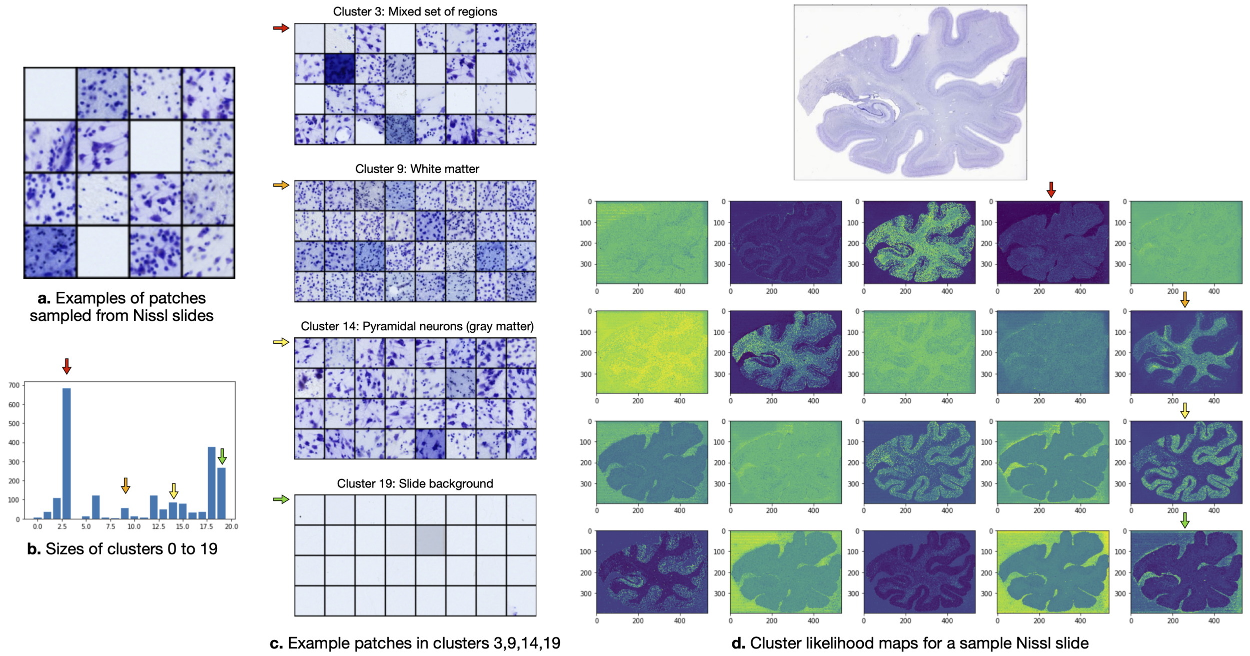

Figure S.6: Illustration of multivariate feature map generation from full-resolution Nissl histology slides. (a). Random examples of 256x256 pixel patches used as input to unsupervised clustering. (b). The sizes of 20 clusters generated by the DeepCluster algorithm Caron et al. [22]. (c). Examples of patches assigned to four selected clusters. Cluster 3 (red arrow) is the largest cluster, contains both foreground and background regions, and appears to be a catch-all cluster with no correspondence to any anatomical area. Cluster 9 contains patches with densely packed oligodendrocytes and no neuronal bodies, corresponding to white matter regions. Cluster 14 contains patches with pyramidal neurons, corresponding to gray matter. Cluster 19 contains slide background patches. (d). Feature maps derived from a single full-resolution Nissl slide. Each feature map in the grid corresponds to a single cluster. Each pixel in each feature map corresponds to a 256x256 patch in the full-resolution slide and quantifies the likelihood that this patch belongs to the given cluster. These feature maps are used to map Nissl slides to lower-resolution images with "MRI-like" appearance to aid registration between MRI and histology. Note: plots (a), (b), (c) are generated from a held out validation set of 2092 patches.

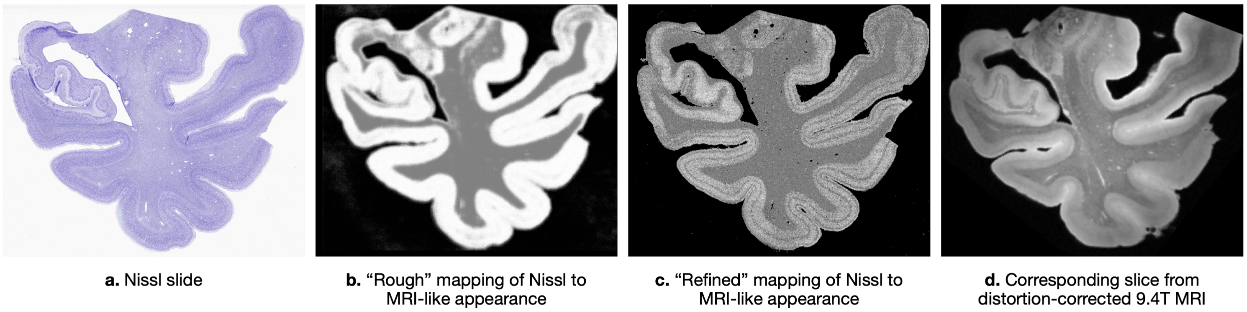

Figure S.7: Intensity mapping from full-resolution Nissl histology slides to MRI-like appearance that facilitates Nissl-to-MRI registration. (a). A sample histology slide. (b). The "rough" MRI-like appearance obtained by classifying the 20-channel feature image (see Figure S.6) into gray matter, white matter, and background regions and taking a weighted sum of the class posterior probability maps. The rough mapping does not require prior registration between Nissl and MRI, and is thus used in the initial registration steps. (c) The "refined" MRI-like appearance obtained by linear fitting of the 20 channels to the MRI intensity after initial Nissl-MRI registration. (d). The MRI slice. The refined MRI-like appearance has fine features, such as the hypointense line in the cortical gray matter, that closely matches that of the MRI . By remapping Nissl to the resolution and appearance similar to that of the MRI, we are able to perform affine and deformable registration using conventional techniques and similarity metrics.

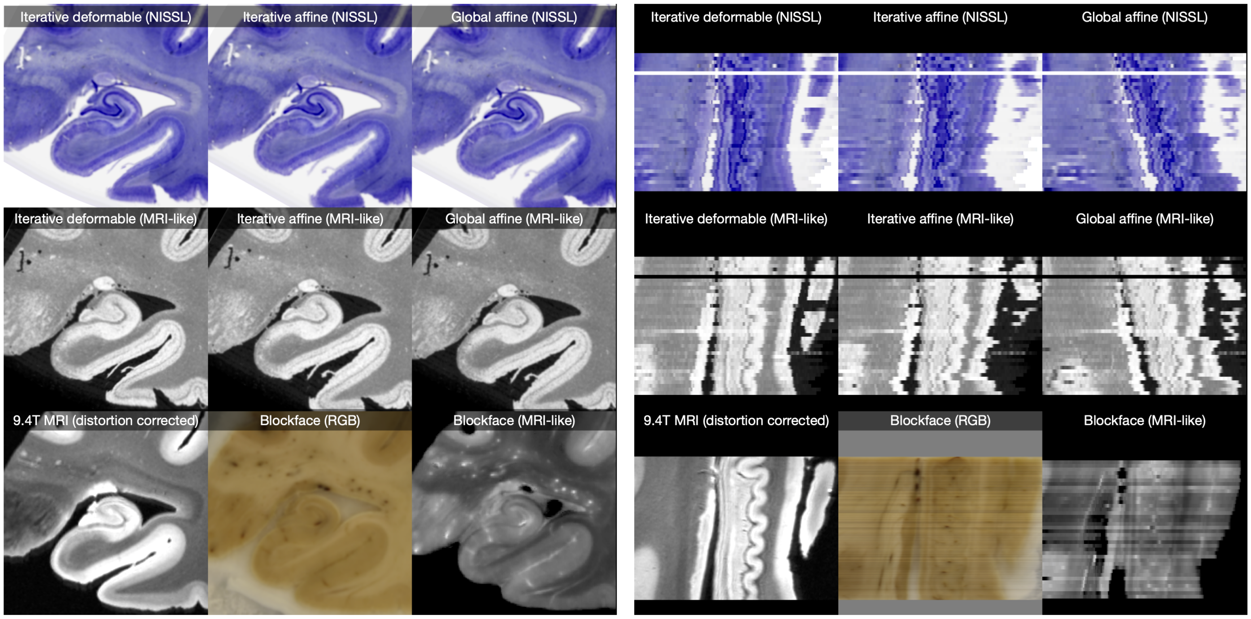

Figure S.8: Example of NISSL to MRI registration for one block covering the hippocampal body. The panel on the left shows a coronal slice, and the panel on the right shows a sagittal slice, on which differences in 3D reconstruction of the histology stack are noticeable. Over the three stages of registration (global affine, iterative affine, iterative deformable), the quality of the 3D reconstruction relative to the 9.4T MRI volume improves. In particular, considerable shape correction can be observed between the global affine result and the iterative affine result in the sagittal view. The blockface image and derived MRI-like volume are also shown.

Table S.2: Anatomical curves used for registration accuracy evaluation.

| **Curve** | **Description** | **Location along the anterior-posterior extent of the MTL** |
|---|---|---|
| C1 | Inner border of the entorhinal cortex up to the top of the collateral sulcus | Anterior tip of the hippocampus |
| C2 | Outer border of the subiculum, pre-subiculum, para-subiculum and entorhinal cortex | Most anterior slice where the hippocampal sulcus opens |
| C3 | Inner border of the stratum radiatum lacunosum moleculare (SRLM) of the hippocampus | Most anterior slice of the hippocampal body (immediately posterior to the posterior tip of the uncus). |
| C4 | Lateral outer border of the hippocampus | 6 mm posterior to the intralimbic gyrus (caudal end of the uncus) |

Table S.3: Demographic and clinical details of the in vivo MRI study participants whose T1-weighted and FLAIR MRI scans were used to generate in vivo brain templates (CN: cognitively normal control; MCI: mild cognitive impairment; AD: probable AD).

| **In Vivo MRI Cohort** | |
|---|---|
| N | 23 |
| Age | 71.4±7.5 |
| Sex | 12 Female, 11 Male |
| Education (Years) | 15.7±3.0 |
| MMSE | 27.9±2.3 |
| Clinical Diagnosis | 11 CN, 10 MCI, 1 AD |

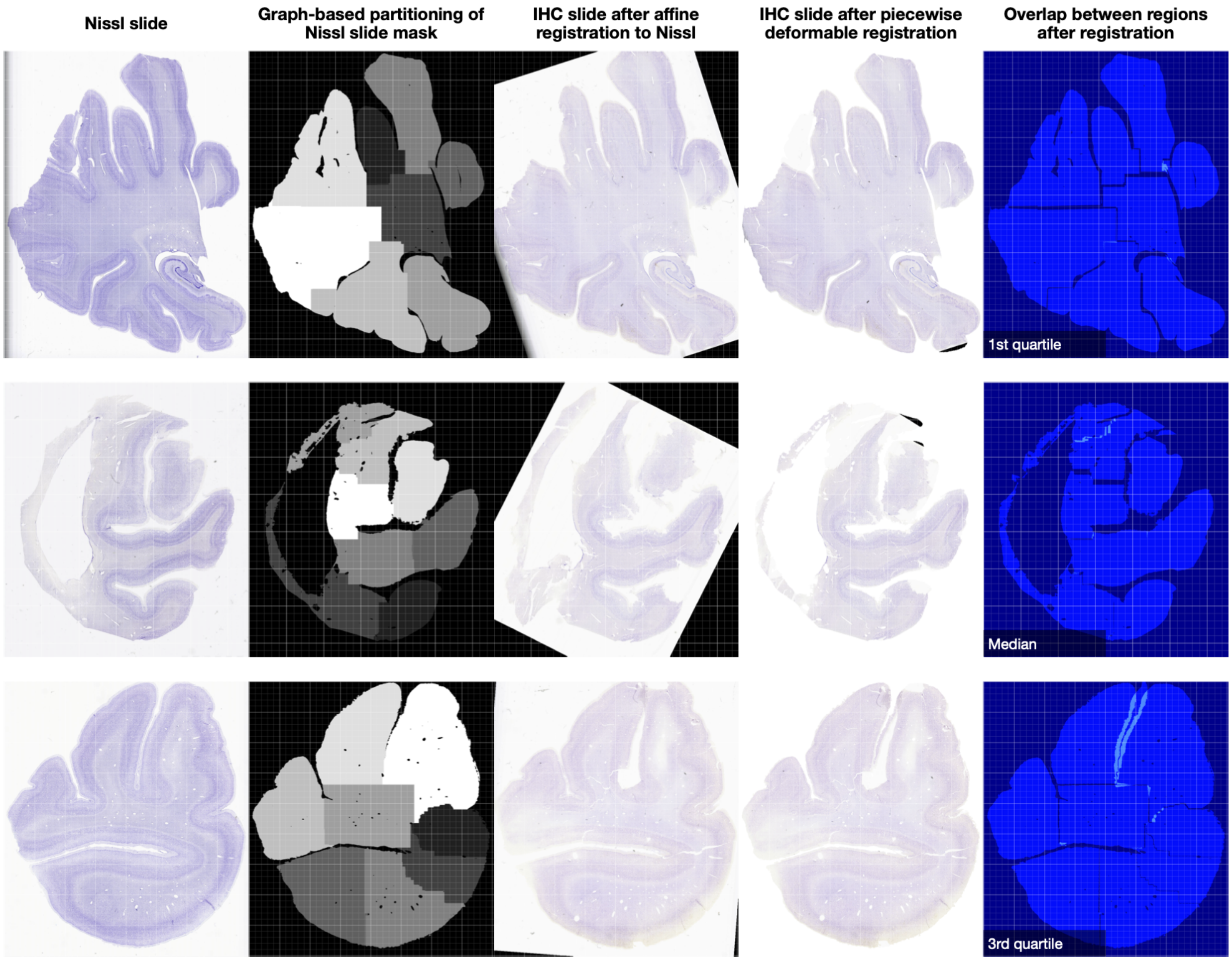

Figure S.9: Examples of piecewise deformable registration between Nissl slides and anti-tau immunohistochemistry (IHC) slides. The foreground mask of the Nissl slide (a) is partitioned into eight regions (b). The IHC slide is first rigidly registered to the Nissl slide (c), and then affine and deformable registration is performed for each region separately (d). The overlap between regions shown in (e) indicates areas of registration inconsistency. The examples shown correspond to the 25th, 50th and 75th percentile of the overlap across all 1224 slides in the study. The piecewise registration approach helps account for tearing of the tissue and large displacements of tissue peninsulas separated by narrow gaps.

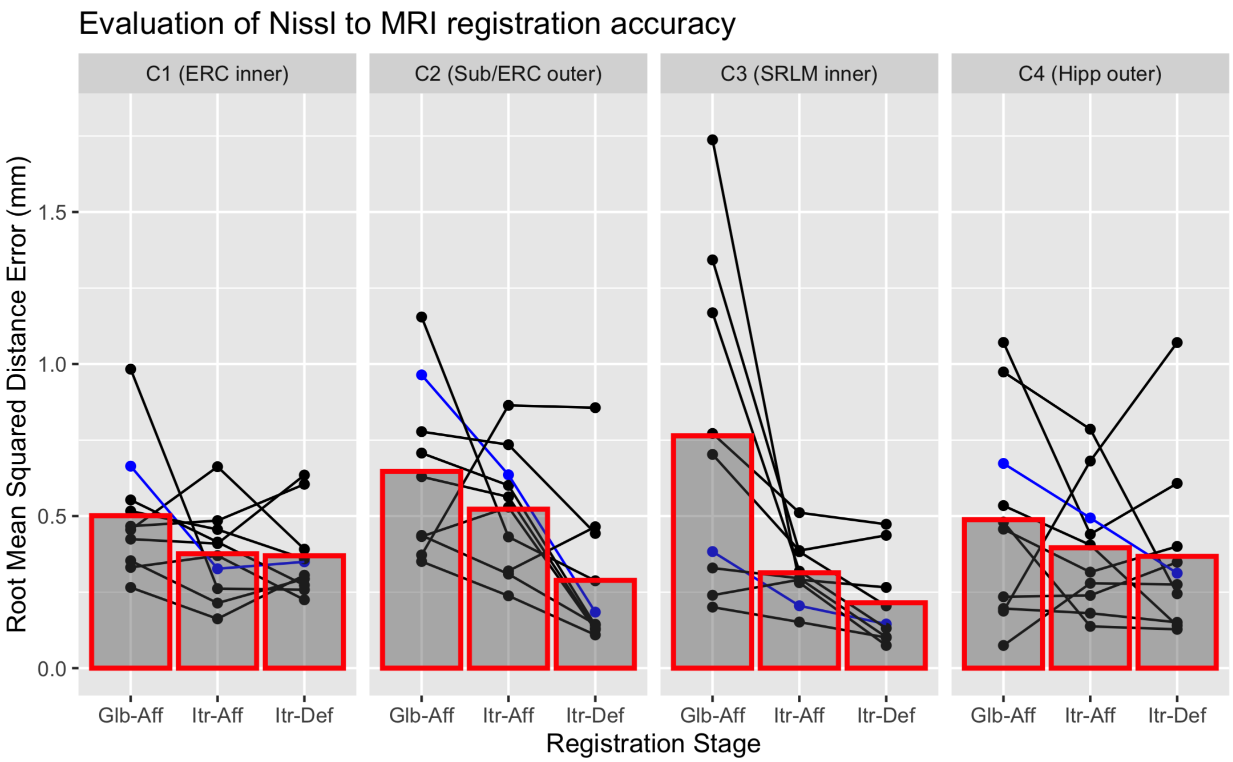

Figure S.10: Analysis of registration accuracy between Nissl and MRI based on anatomical curves traced manually in both modalities. The plot reports curve mismatch in terms of root mean square distance between the curves traced in Nissl and MRI. The four columns correspond to four different anatomical curves listed in Table S.2. Within each subplot, the red bars plot the mean curve mismatch for three registration stages: global affine (Glb-Aff), iterative affine (Itr-Aff) and iterative deformable (Itr-Def). Dots connected by line segments represent individual specimens. Registration for four cases labeled in blue (median performance for each curve at the Itr-Def stage) are visualized in Supplemental Figure S.11.

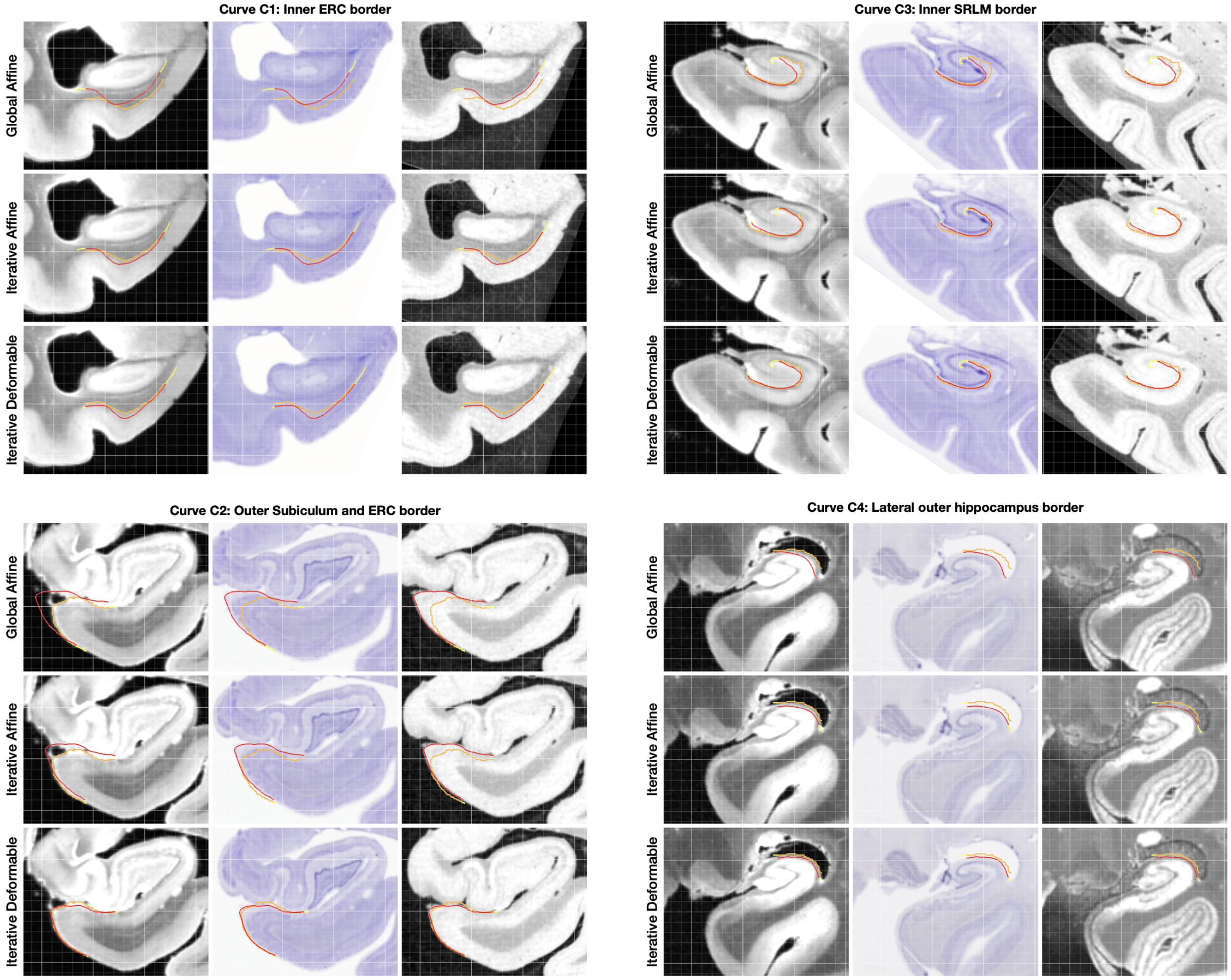

Figure S.11: Examples of curve-based evaluation of registration between 9.4 Tesla MRI and Nissl slides. For each of the four anatomical curves (defined in Supplemental Table S.2), registration between an MRI slice and corresponding Nissl histology slice is shown at three stages: global affine registration, iterative affine registration, and iterative deformable registration. In each set of examples, the MRI slice is shown on the left, the Nissl slice in the middle, and the "MRI-like" image derived from the Nissl slide using intensity remapping is shown on the right. The registration evaluation metric plotted in Figure S.10 measures the root mean square distance between the red curve (traced in MRI space) and orange curve (traced independently in Nissl space). Portions of curves excluded from the distance computation are shown in yellow. The four examples shown here correspond to median registration performance for each curve, and are identified by blue lines in Figure S.10 .

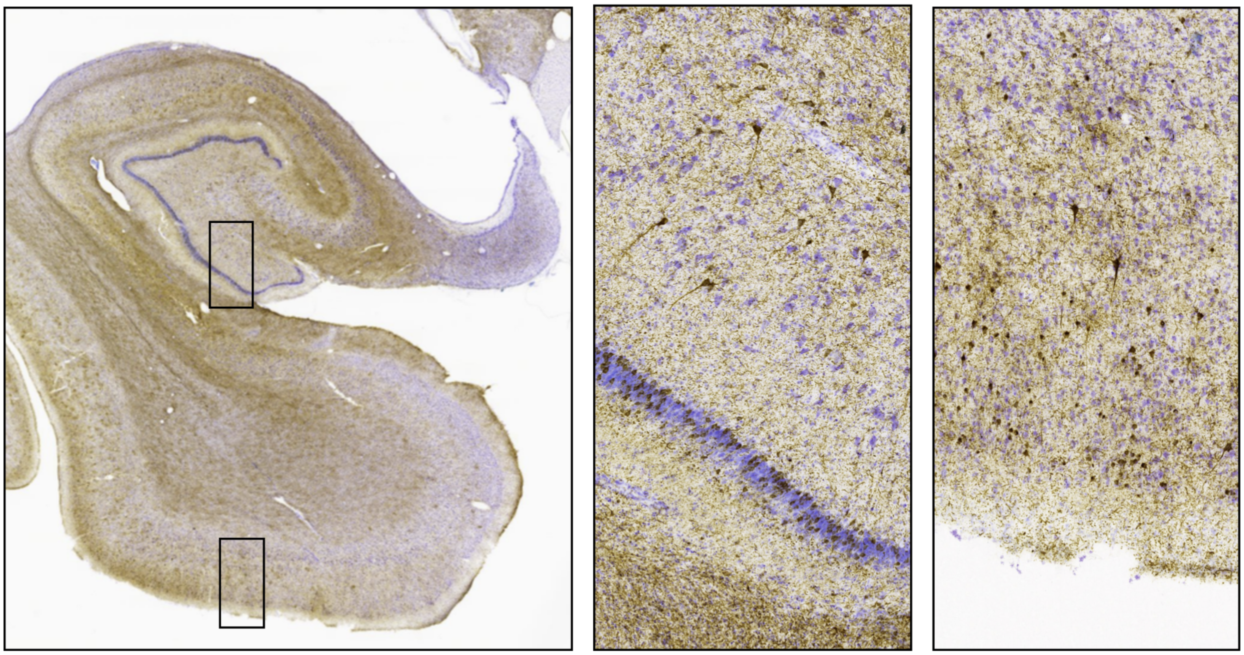

Figure S.12: A sample anti-tau immunohistochemistry slide from a specimen with corticobasal degeneration pathology that was rated Braak stage B0 based on the staining of the collateral hemisphere with the GT38 stain, which differentiates AD-like NFT pathology from FTLD-tau [32, 33]. Due to the visual similarity of intracellular tau inclusions in this case to AD-like NFTs, the NFT burden map generated by our pipeline in this case (Case 16 in Figure 5) exhibits high levels of NFT burden.

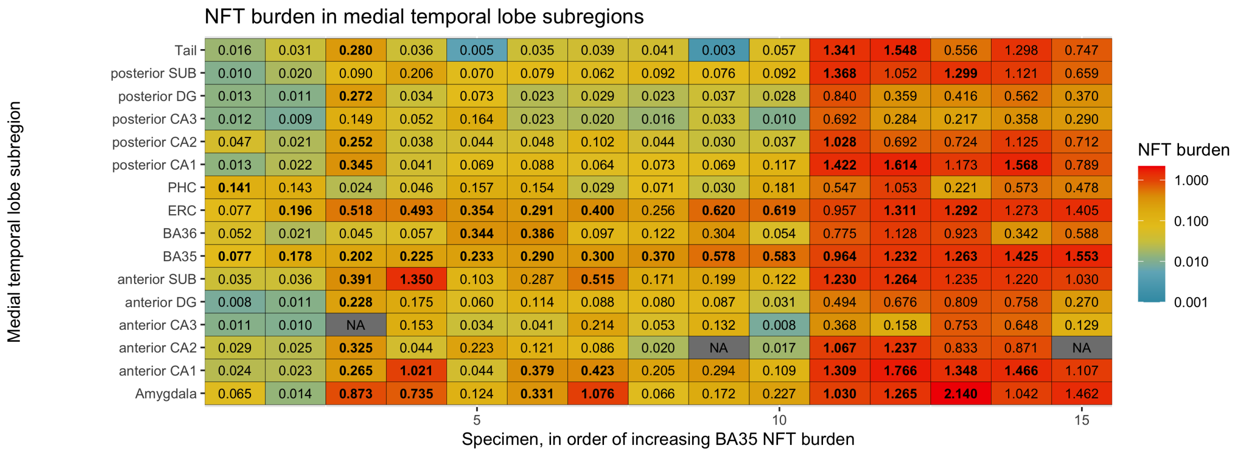

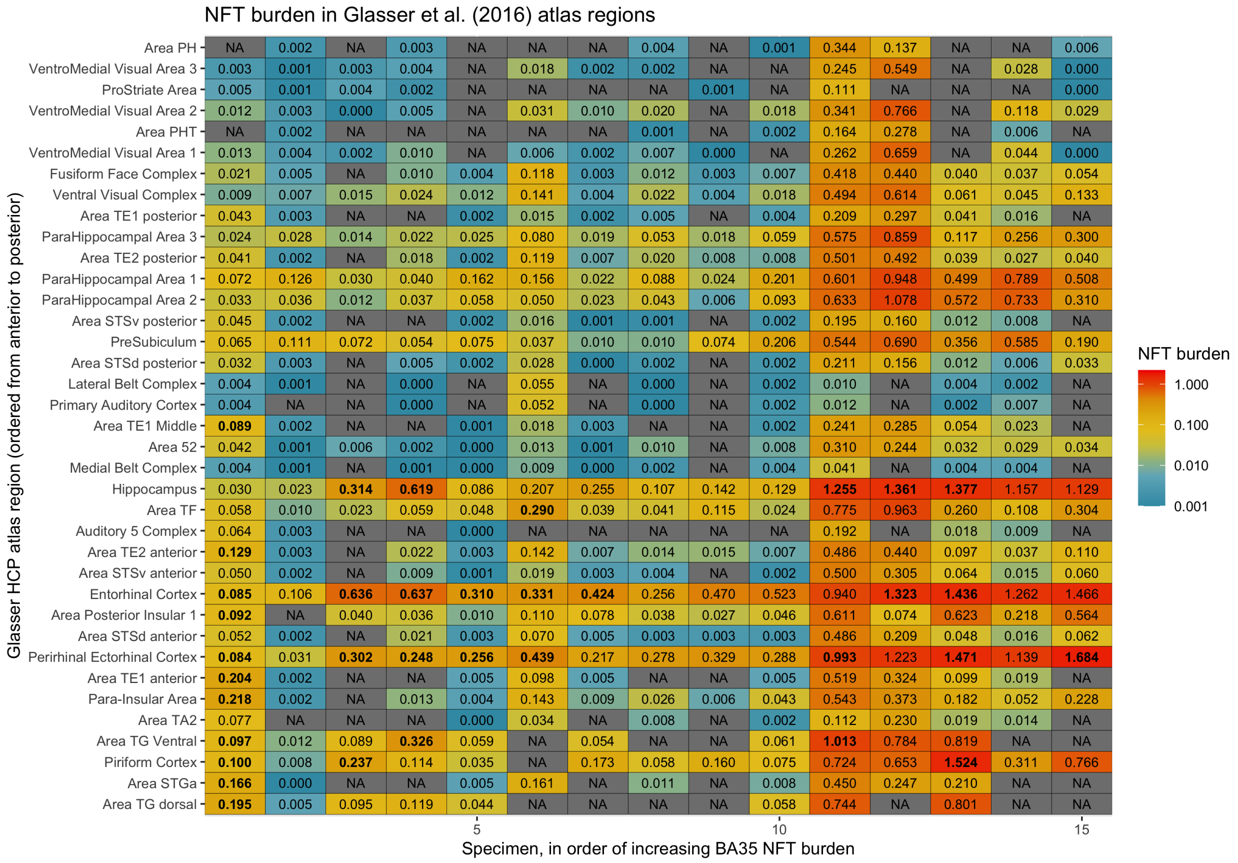

Figure S.13: Summary measures of tau neurofibrillary tangle (NFT) burden in anatomical regions of interest for the 15 specimens included in group analysis. The summary measure is the 90th percentile of the voxel-wise NFT burden measure over each region in each specimen. The top plot shows medial temporal lobe anatomical regions, and the bottom plot shows anatomical regions from the Glasser et al. [34] Human Connectome Project atlas. Specimens are arranged from left to right in the order of the Brodmann Area 35 (BA35) summary NFT burden measure, and regions for which the NFT burden measure is equal or greater to BA35 NFT burden are marked with bold text. These measures suggest that even when BA35 NFT burden is mild (0.25 or less), NFT pathology is frequently found in other regions. Cells labeled "NA" indicate no available measurement due to lack of anti-tau stained IHC overlapping the region.

Table S.4: Registration parameters used in different stages of the histology reconstruction pipeline. Registration steps labeled A-F in Supplemental Figure S.4 are broken down into stages, and for each stage, details of registration are provided. Registration direction describes which images play the roles of fixed image and moving image in the registration, and whether the main output of the registration uses the forward transformation (mapping fixed image coordinate space to moving image coordinate space) or the inverse transformation. Initialization describes how the initial transformation parameters are derived. Masking describes whether a binary image is used as a fixed image mask (limiting the computation of the similarity metric and its gradient to a subset of voxels) or moving image mask (limiting the set of voxels in the moving image that are sampled for similarity metric computation). Multi-resolution schedule describes the maximum number of iterations of registration performed at successive multi-resolution levels, e.g., 100x40x10 signifies 100 iterations at 4x downsampling, 40 iterations at 2x downsampling, and 10 iterations at full resolution. Similarity metric describes the image similarity metric used. The notation NCC 2x2x2 signifies that the normalized cross-correlation metric is used with patch radius of 2 voxels (i.e., patch size 5x5x5 voxels). Regularization describes the amount of "fluid-like" and "diffusion-like" regularization [74] used during deformable registration.

| **A. 9.4T MRI aligned and warped to match the 7T MRI (Section 5.5.1)** | | |
|---|---|---|
| **Stages** | Stage I | Stage II |
| **Description** | Affine registration | Deformable registration |
| **Tool** | ITK-SNAP | GreedyReg |
| **Registration Direction** | 7T MRI as fixed, 9.4T MRI as moving | 9.4T MRI as fixed, 7T MRI as moving, output inverted |
| **Initialization** | Manual rotation and translation in ITK-SNAP | Inverted transformation from Stage I |
| **Masking** | None | 9.4T MRI foreground region used as fixed mask in cases when 9.4T has partial coverage |
| **Multi-resolution schedule** | ITK-SNAP default | 100x100x100x40 |
| **Similarity metric** | ITK-SNAP default | NCC 8x8x8 |
| **Regularization** | | (3mm, 0.2mm) |

| **B. Distortion-corrected 9.4 MRI aligned to the 3D block face volume (Section 5.5.2)** | | | |
|---|---|---|---|
| **Stages** | Stage I | Stage II | Stage III |
| **Description** | Rigid registration | Affine registration | Affine registration |
| **Tool** | GreedyReg | GreedyReg | GreedyReg |
| **Registration Direction** | 7T MRI as fixed, MRI-like block face volume as moving | 7T MRI as fixed, MRI-like block face volume as moving | MRI-like block face volume as fixed, 9.4T MRI as moving |

| | | | |
|---|---|---|---|
| **Initialization** | Manual placement of block face stacks relative to 7T MRI in ITK-SNAP (based on 3D printed molds) | Stage I | Inverted transformation from Stage II |
| **Masking** | 7T MRI foreground mask used as fixed mask | 7T MRI foreground mask used as fixed mask | Block face foreground segmentation (from random forest) used as fixed mask |
| **Multi-resolution schedule** | 60x40x0 | 60x40x0 | 60x40x0 |
| **Similarity metric** | NCC 4x4x4 | NCC 4x4x4 | NCC 4x4x4 |
| **Regularization** | | | |

| **C. Tau IHC slides aligned and warped to adjacent NISSL slides (Section 5.5.5)** | | | |
|---|---|---|---|
| **Stages** | Stage I | Stage II | Stage III |
| **Description** | Rigid registration on the whole slide | Rigid registration performed separately for 8 "pieces" | Deformable registration performed separately for 8 "pieces" |
| **Tool** | GreedyReg | GreedyReg | GreedyReg |
| **Registration Direction** | Nissl as fixed, IHC as moving | Nissl as fixed, IHC as moving | Nissl as fixed, IHC as moving |
| **Initialization** | Brute force search in GreedyReg (10,000 iterations, random sampling of orientation and flips) | Stage I | Stage II |
| **Masking** | Nissl slide foreground region (from random forest) used as fixed mask | Nissl slide foreground region partitioned into 8 "pieces", each used as fixed image mask | Same as Stage II |
| **Multi-resolution schedule** | 100x40x10x0 | 100x40x10x0 | 100x40x10x0 |
| **Similarity metric** | NCC 16x16 | NCC 16x16 | NCC 16x16 |
| **Regularization** | | | (3.0mm, 0.5mm) |

| **D, E: MRI-like Nissl slides stacked in 3D and aligned to the 9.4T MRI (Section 5.5.4)** | | | | |
|---|---|---|---|---|
| **Stages** | Stage I | Stage II | Stage III | Stage IV |
| **Description** | Nissl slides stacked in 3D by affine registration of each slide to adjacent slides and composing transformations using a graph-based strategy | Each Nissl slide registered to corresponding slice of 9.4T MRI (in block face space). The single transformation that achieves best metric between all MRI slices and the Nissl stack is selected as output. Uses "rough" MRI-like Nissl appearance (Supplemental Figure S.7). | Iterative affine registration between each Nissl slide, two adjacent Nissl slides, and corresponding MRI slide. Uses "rough" MRI-like Nissl appearance. | Iterative deformable registration between each Nissl slide, two adjacent Nissl slides, and corresponding MRI slide. Uses "refined" MRI-like Nissl appearance. |
| **Tool** | stack_greedy | stack_greedy | stack_greedy | stack_greedy |

| | | | | |
|---|---|---|---|---|
| **Registration Direction** | For each Nissl slide, it is used as the fixed image, nearby Nissl slides (up to 1.6mm away) used as moving | MRI slice as fixed, Nissl slide as moving | Nissl slide as fixed, adjacent Nissl slides and MRI slice as moving | Same as Stage III |
| **Initialization** | Brute force search in stack_greedy (4,000 iterations, random sampling of orientation and flips) | Brute force search in stack_greedy (4,000 iterations, random sampling of orientation and flips) | Stage II | Stage III |
| **Masking** | Nissl foreground region (from random forest) used as fixed mask | | Same as Stage I | Same as Stage I |
| **Multi-resolution schedule** | 100x40x0 | 100x40x10 | 100x40x10 (for 10 iterations) | 40x80x80 (for 10 iterations) |
| **Similarity metric** | NCC 8x8 | NCC 8x8 | NCC 8x8 | NCC 8x8 |
| **Regularization** | | | | (3.0mm, 0.25mm) |

| **F: Group Template Generation (Section 5.5.8)** | | | | |
|---|---|---|---|---|
| **Stages** | Stage I | Stage II | Stage III | Stage IV |
| **Description** | Construct unbiased population template from in vivo T1-MRI using one iteration of affine registration and five iterations of deformable registration. | Affine registration between 9.4T MRI and the in vivo FLAIR template. | Deformable registration between 9.4T MRI and the in vivo FLAIR template. | Construct unbiased population template using 9.4T MRI aligned to the in vivo FLAIR template using 10 iterations of deformable registration. |
| **Tool** | GreedyReg | GreedyReg | GreedyReg | GreedyReg |
| **Registration Direction** | Iteratively updated average image as fixed, individual in vivo T1-MRI scans as moving. | 9.4T MRI downsampled to $0.5 \times 0.5 \times 0.5\,\mathrm{mm}^3$ resolution as fixed, FLAIR template as moving | Same as Stage II. Inverse transformation is used to map 9.4T MRI to template space. | Iteratively updated average image as fixed, individual scans as moving. |
| **Initialization** | | Manual rotation/translation in ITK-SNAP | Stage II | Stage III |
| **Masking** | | MRI foreground as fixed mask | Same as Stage II | Iteratively updated template mask as fixed mask, MRI foreground as moving mask |
| **Multi-resolution schedule** | 60x20x0 (affine), 60x20x0 (deformable) | 100x40x0 | 100x100x40 | 100x40x20 |
| **Similarity metric** | NCC 2x2x2 | NCC 2x2x2 | NCC 2x2x2 | NCC 2x2x2 |
| **Regularization** | (3.0mm, 0.5mm) | | (3.0mm, 1.0mm) | (3mm 0.5mm) |
